# Local nucleolar surveillance mechanisms maintain rDNA stability

**DOI:** 10.1101/2025.01.20.633984

**Authors:** Ruofei Liu, Jiachen Xuan, Junqi Pan, Shalini Sundramurthi Chelliah, Rhynelle Dmello, Karla J. Cowley, Shannon Mendez, Yangyi Zhang, Kezia Gitareja, Matthew Wakefield, Andrew Deans, Clare Scott, Keefe T. Chan, Kaylene J. Simpson, Jian Kang, Elaine Sanij

## Abstract

The nucleoli are subdomains of the nucleus that form around actively transcribed ribosomal RNA (rRNA) genes. The highly repetitive and transcribed nature of the rRNA genes (rDNA) by RNA polymerase I (Pol I) poses a challenge for DNA repair and replication machineries. Here, we profile the nucleolar proteome and the chromatin landscape of stalled replication sites upon rDNA damage to characterize the early steps of nucleolar DNA damage response (nDDR). We observed early dynamics in nucleolar-nucleoplasmic proteome localization and identified nucleolar replication stress signatures involving chromatin remodeling networks, transcription-replication conflicts and DNA repair. Our findings define localized surveillance mechanisms that activate the nDDR. Further, we identified that upon rDNA damage, nucleolar RNA Polymerase (Pol) II binds to intergenic rDNA sequences and generates R-loops (DNA:RNA hybrid structures) that are essential for recruiting nDDR factors. Using a boutique CRISPR-Cas9 synthetic lethal screen of DNA repair factors with inhibitors of RNA Pol I transcription, we identified an unexpected protective role for the DNA translocase RAD54L in nDDR. Loss of Rad54L increases nucleolar R-loops and rDNA damage leading to defects in nucleolar structure and enhanced sensitivity to PARP and RNA Pol I inhibitors. Altogether, our study uncovers localized surveillance networks within the nucleolus that respond to rDNA damage. These insights expand our understanding of the molecular mechanisms governing nDDR and opens new avenues for developing nDDR-targeting therapies.

## Introduction

RNA polymerase I (Pol I) transcription of rDNA constitutes more than 60% of all cellular transcription^1^. The rDNA repeats are organized in clusters of 70-80 copies on the short arm of the five acrocentric chromosomes in human cells. RNA Pol I transcribes a subset of the rDNA repeats to produce the 47S precursor-rRNA (pre-rRNA) within the nucleoli, which serve as the sites of RNA Pol I transcription and ribosomal subunit assembly. The rDNA repeats are intrinsically unstable due to their high transcriptional activity and repetitive nature, making them susceptible to DNA damage and frequent rearrangements across the genome. The instability of the rDNA loci is increased upon loss of genome maintenance factors such as BLM and WRN, emphasizing the need for surveillance of rDNA loci integrity^2^.

Several unique characteristics of the rDNA repeats, including their repetitive nature, high Pol I transcription rates and high GC content, present risks of genomic instability^3^. The head-to-tail arrangement of rDNA copies per locus allows intrachromosomal and interchromosomal recombination events, which can lead to alterations in rDNA copy number and genomic rearrangements^4,5^. Furthermore, high levels of transcription render the rDNA clusters vulnerable to transcription-replication conflicts (TRCs). A head-on collision between the transcription machinery and DNA replication forks triggers local accumulation of positive DNA supercoiling, replication fork stalling, torsional stress and the accumulation of RNA:DNA hybrids (R-loops)^6^, which leads to double strand breaks (DSBs). rDNA DSBs are repaired by the homologous recombination (HR), non-homologous end-joining (NHEJ) repair and single-strand annealing pathways that are activated depending on the cell cycle stage, the extent of rDNA damage and persistence of DSBs^5,7^. As such, high fidelity in rDNA repair is critical for maintaining genome integrity.

The transit from low to high levels of rDNA damage is associated with ATM-mediated inhibition of RNA Pol I transcription and large-scale reorganization of nucleolar architecture that involves localization of rDNA DSBs to nucleolar caps at the periphery. It has been suggested this restructuring of the nucleolus serve as a mechanism to separate rDNA from different chromosomes to prevent detrimental recombination events^8^. This transition is also associated with a shift from NHEJ to HR type of repair. In agreement with this, HR factors were shown to be recruited to nucleolar caps after rDNA damage^8,9^, presumably to restrict HR activity to damaged rDNA at the nucleolar periphery.

While a vast majority of DDR factors involved in DNA damage sensing, signalling and repair are reported to reside within the nucleoli, most studies have focussed on assessing the mobilization of specific DDR factors to the nucleoli upon rDNA damage. For instance, the ATM-activated nucleolar protein Treacle (also known as TCOF1) is known to recruit the MRN complex and the DNA topoisomerase II binding protein 1 (TOPBP1) to rDNA DSBs in the nucleolus^10,11^. Thus far, our understanding of the dynamics of early DDR upon rDNA damage and the mechanisms that secure the integrity of the rDNA loci has been limited to assessing the association of specific DDR factors with the nucleoli.

Here, we employed an inducible rDNA damage model to assess changes in the nucleolar proteome and to profile the chromatin landscape at stalled replication sites upon rDNA damage. We have identified signatures of replication stress at the rDNA loci and local nucleolar surveillance networks that enable a dynamic DDR to DSBs at rDNA. Moreover, we have uncovered a new role for nucleolar RNA Pol II in binding intergenic spacer (IGS) rDNA regions to produce anti-sense IGS rRNAs and generating R-loops to promote nDDR.

In addition, we utilized a boutique CRISPR-Cas9 screen to assess the interactions of DNA repair pathways with RNA Pol I inhibitors and identified a unique synthetic lethal interaction with the DNA translocase RAD54L. RAD54L plays distinct roles in regulating RAD51 during HR repair and in restraining replication fork speed. Here, we uncover that nucleolar RAD54L plays critical roles in resolving replication stress, initiating DDR signaling and protecting the fidelity of the rDNA loci. Our data highlights an important protective role for RAD54L in resolving R-loops at the rDNA loci and that the loss of RAD54L leads to increased rDNA damage and sensitivity to PARP and Pol I inhibitors. Altogether, our data demonstrate dynamic local nucleolar networks and mechanisms that responds to rDNA damage and nucleolar replication stress to secure the integrity of the rDNA loci.

## Results

### rDNA damage induces nDDR and G2/M cell cycle arrest

Gene editing tools, especially the CRISPR-Cas9 system, allows targeted induction of DSBs to almost any locus of the genome in a controllable manner to study specialized DDR pathways associated with specific nuclear compartments such as the nucleoli^12^. To gain insights into the nature of nDDR, we established a cell line model of two clonal OVCAR8 ovarian cancer cell lines stably expressing doxycycline-inducible endonuclease Cas9 and two small guide RNAs (sgRNAs) targeting the rDNA (sgRNA-rDNA), named GRASP (Guide RNA for rDNA Specificity) (Fig 1A). Treatment with doxycycline for 24 hours (h) induced Cas9 and pCHK2 T68, a marker of active DDR (Figure S1A). The induction of Cas9 also correlated with the detection of the DNA damage marker γH2AX at the nucleolar periphery (Fig 1B).

**Figure 1.**
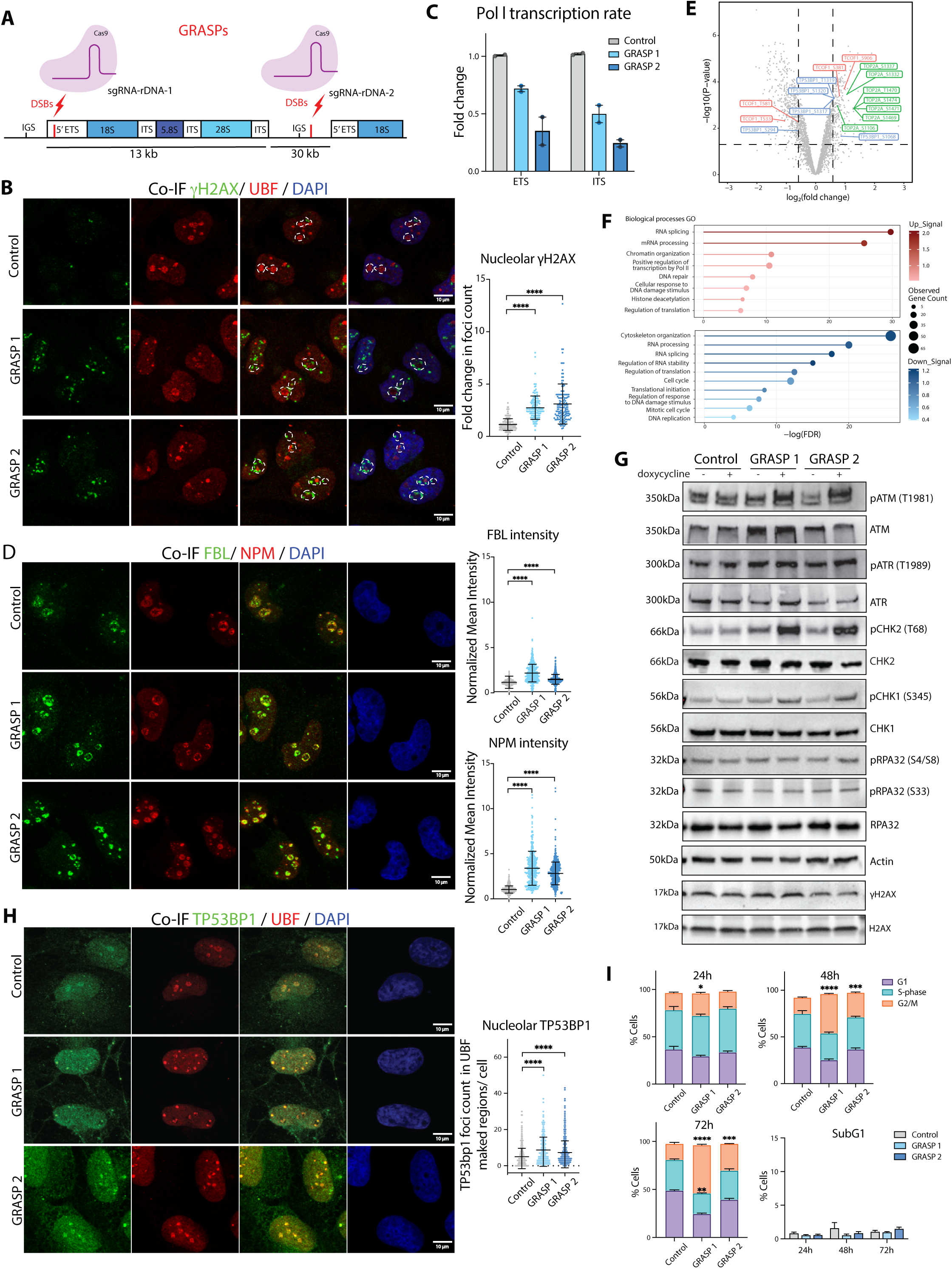
rDNA damage induces nucleolar DDR and G2/M cell cycle arrest. **A**. Schematic illustrating the rDNA damage model and the location of sequences targeted by sgRNA-rDNA. **B**. Co-immunofluorescence (Co-IF) analysis of γH2AX and UBF, a marker of active rDNA^46^, in cells treated with doxycycline (100 ng/ml) for 24h. Representative images from three independent biological experiments. Nucleolar γH2AX foci were quantitated using ImageJ. A total of 300 cells per treatment were analyzed across three experiments. Error bars represent mean ± SD. Statistical analysis was conducted using an unpaired t-test; **** is p-value ≤ 0.0001. **C**. Pol I transcription level determined by quantitative real-time PCR (qRT-PCR) using primers specific to internal and external transcribed spacer (ITS and ETS) sequences of the 47S rRNA precursor (Table S12). Expression levels were normalized to Vimentin mRNA and expressed as fold changes relative to control OVCAR8 cells treated with doxycycline (100 ng/ml) for 24h. Data represent means from two independent biological replicates. Error bars indicate mean ± SEM. **D**. Co-IF analysis of fibrillarin (FBL) and nucleophosmin (NPM) in cells treated with doxycycline (100 ng/ml) for 24h. Representative images from three independent experiments. Signal intensity for FBL and NPM was quantified using CellProfiler and normalized to the median value of control cells per experiment. Over 400 cells were analyzed per condition across three independent experiments. Error bars represent mean ± SD. Statistical analysis was conducted using an unpaired t-test; **** is p-value ≤ 0.0001. **E**. Volcano plot representing the differential abundance of phosphoproteins between the GRASP 1 and control cells following treatment with doxycycline (100 ng/ml) for 24h. Blue, red, green dots indicate up-regulated levels of phosphorylated forms of TP53BP1, TCOF1 and TOP2A (log_2_ fold change ≥ 1; p-value < 0.05), respectively. **F**. Lollipop plot of gene ontology (GO) enrichment analysis of altered phosphoproteins in GRASP cells, with up-regulated biological processes shown in red and down-regulated processes in blue. **G**. Western blot analysis of key DDR proteins in cells treated 100 ng/ml doxycycline for 24h. Representative of three independent biological replicates. **H**. Co-IF analysis of TP53BP1 and UBF in GRASP and control cells treated with doxycycline (100 ng/ml) for 24h. Representative images of three biologically independent experiments. Quantification of foci was performed with CellProfiler on over 300 cells per condition across three experiments. Error bars represent mean ± SD. Statistical analysis was performed using an unpaired t-test; **** p-value ≤ 0.0001. **I**. Analytical cell cycle analysis of BrdU incorporation as a function of DNA content using FACS. Cells were treated with doxycycline (100 ng/ml) for 24h, 48h and 72h, and labelled with BrdU for 30 minutes prior to harvest. Histogram plots display the percentage of G1 and G2/M and S-phase BrdU-labelled cells, and SubG1 cell populations (gating strategy shown in Fig. S1E). Error bars represent mean ± SEM. Statistical analysis was performed using unpaired t test; * p-value ≤ 0.05, ** p-value ≤ 0.01, *** p-value ≤ 0.001, **** p-value ≤ 0.0001.

DSBs in rDNA can lead to repression of RNA Pol I transcription and nucleolar segregation^8,13^. As expected, the induction of DSBs at rDNA led to reduced RNA Pol I transcription rate (Fig 1C) and altered nucleolar organization indicated by the localization of nucleolar proteins nucleophosmin (NPM) and fibrillarin (FBL) into dense rings at the nucleolar caps (Fig 1D). Next, we performed proteomic and phospho-proteomic analyses to characterize the molecular response to rDNA DSBs. Interestingly, we did not observe major changes in global protein levels within 24h of inducing rDNA damage (Fig S1B, Table S1). In contrast, phospho-proteomics analysis identified 767 phospho-sites that were differentially regulated in GRASP1 cells compared to control OVCAR8 cells (Fig 1E, Fig S1C and Table S2). Gene ontology analysis of deregulated phospho-proteins in GRASP cells identified an enrichment of factors involved in RNA processing and regulation of RNA polymerase II-mediated transcription but down-regulation of processes related to cell cycle progression, DNA replication and RNA splicing (Fig 1F). The upstream kinase activity inferred from the substrate phosphorylation data using KEA3 and Phosphomatics identified ATM, CHK2 and CDKs signaling activities to be upregulated upon the induction of DSBs in rDNA (Fig S1D). In agreement with this, increased levels of ATM phosphorylation at T1981 were detected in GRASP cells compared to a modest increase in phosphorylation of ATR (Fig 1G). A significant increase in pCHK2 T68 levels and a mild increase in pCHK1 (S345) were observed, indicating that the induction of rDNA DSBs in GRASP cells mainly activates canonical ATM signaling (Fig 1G). Interestingly, total levels of γH2AX were unaltered in GRASP cells suggesting that the DNA damage is localized to the nucleoli as seen in the co-IF analysis (Fig 1B). Similarly, global levels of markers of replication stress, including phosphorylation levels of replication protein A pRPA32 (S33) and (S4/S8) were also unaltered (Fig 1G), indicating limited effects on global replication within 24h of doxycycline treatment of GRASP cells.

Within the top differentially regulated phospho-peptides, phosphorylation of TCOF1 at (S381 and S906) was significantly upregulated upon rDNA DSBs induction (Fig 1E). ATM is involved in phosphorylating TCOF1, a central coordinator of nDDR by recruiting and retaining DDR factors in the nucleolus ^14^. Altered phosphorylation of TOP2A and TP53BP1 was also detected in GRASP cells (Fig 1E), suggesting the presence of replication stress and activation of NHEJ DNA repair, respectively. Indeed, TP53BP1 foci were detected at the nucleoli following the induction of rDNA damage (Fig 1H), indicating NHEJ pathway activation. The induction of rDNA DSBs also impacted cell cycle progression, with GRASP1 cells exhibiting S-phase delay within 24h and both GRASP cell lines exhibiting G2/M arrest within 48-72h upon the induction of rDNA DSBs (Fig 1I and Fig S1E). These findings suggest that sustained nDDR signaling activates NHEJ and cell cycle checkpoints to impede global DNA replication and cell cycle progression.

To define the impact of rDNA damage and activation of nDDR on nucleolar-nucleoplasmic proteome distribution, we performed nucleolar fractionation followed by western blotting (Fig 2A) and mass spectrometry (MS) analysis. Western blot analysis of FBL and the nucleoplasmic protein CIAPIN1 demonstrates the purity of isolated nucleolar and nucleoplasmic fractions (Fig 2A). Furthermore, increases in γH2AX and in pRPA32 (S33) were observed only in the nucleolar fractions upon the induction of rDNA damage (Fig 2B). We also evaluated the fractionation quality by assessing the raw MS data for FBL (Fig S2A) by calculating the percentage of FBL abundance in each compartment. As expected, only 3.47% of total FBL was detected in the nucleoplasm, with approximately 96.53% of FBL residing in the nucleolus. Proteomic data analysis of 3 biological replicates showed that 1,605 proteins were identified in the nucleolus (Fig S2B and Table S3). Gene ontology analysis using STRING GO of the biological processes that are enriched with nucleolar proteins revealed that approximately 13% were involved in ribosome biogenesis, 8.8% in RNA splicing, 6.9% in chromatin organization, and around 13.6% in DDR and cell cycle regulation (Fig 2C and Table S4). These findings are consistent with previous reports depicting the nucleolus as a central hub that coordinate essential biological processes crucial for cell growth in response to stress^15^.

**Figure 2.**
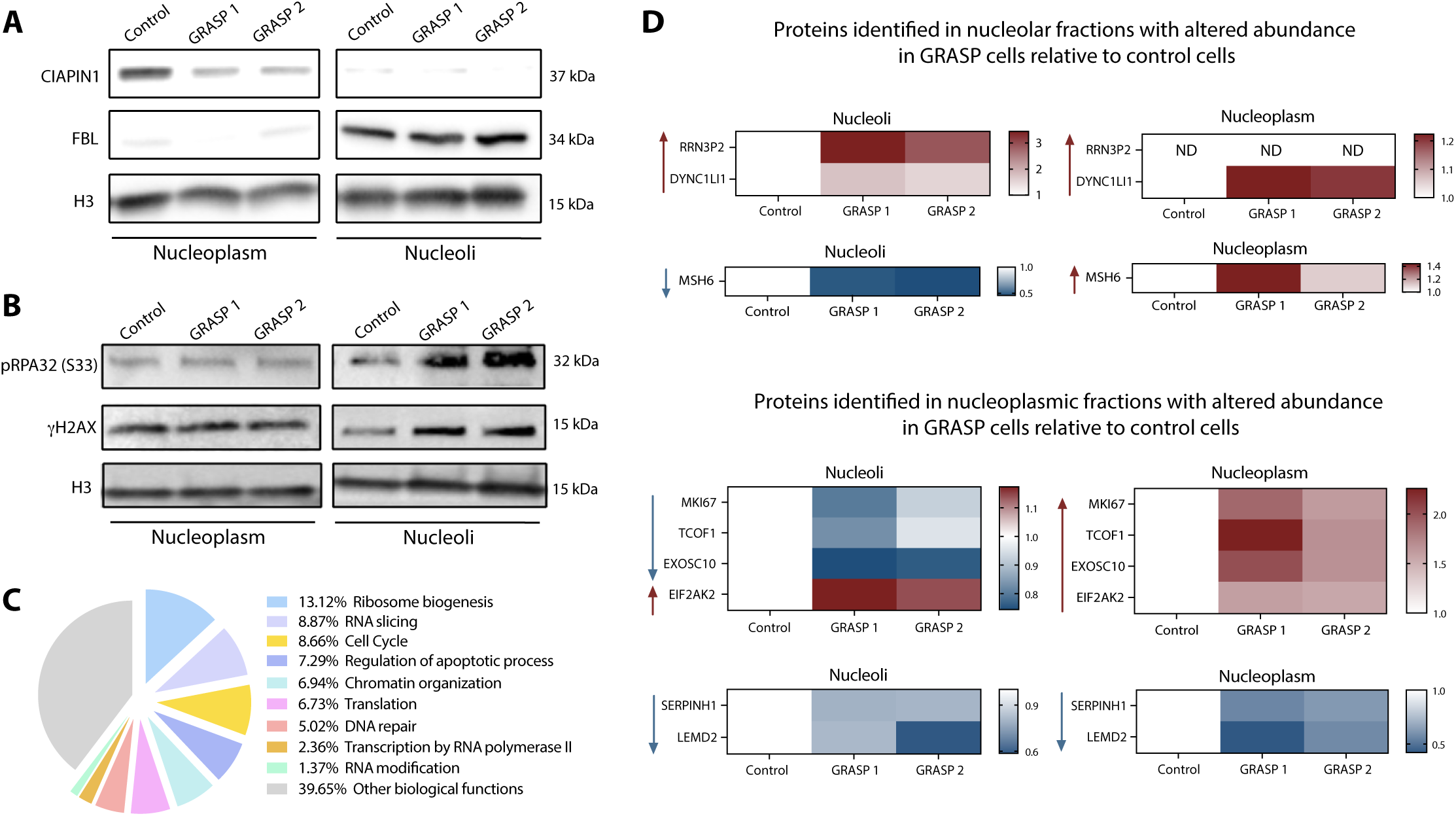
Assessment of subnuclear proteome composition upon rDNA damage. **A**. Western blot analysis of the nucleolar marker FBL and nucleoplasmic marker cytokine-induced apoptosis inhibitor 1 (CIAPIN1) of nucleolar and nucleoplasmic fractions from control OVCAR8 and GRASP cells treated with 100 ng/ml doxycycline for 24h. Histone H3 was used as a loading control. **B**. Western blot analysis of pRPA32 (S33) and γH2AX levels in nucleolar and nucleoplasmic fractions from control and GRASP cells treated with 100 ng/ml doxycycline for 24h. Histone H3 was used as a loading control. **C**. Pie chart representing STRING GO network analysis of the biological processes enriched in the 1,605 proteins that localized to the nucleolus of control OVCAR8 cells. **D**. MS analysis of nucleolar and nucleoplasm fractions following the induction of DSBs at rDNA. Three biological replicates were prepared for the nucleolus and nucleoplasm of control and the two GRASP cell lines treated with 100 ng/ml doxycycline for 24h. The workflow for MS data analysis is presented in Fig S2B. Heatmap illustrating the fold change in abundance of enriched and depleted proteins in the nucleolus and nucleoplasm in GRASP cells compared to control cells. ND indicated not detected in the subnuclear proteome.

We then compare the changes of protein abundance in nucleolar and nucleoplasmic fractions in the two GRASP cell lines relative to control cells and identified only 3 factors that exhibited either increased or reduced abundance in the nucleoli following doxycycline treatment for 24h (log2 mean difference ≥ 0.58 or ≤ –0.58 equivalent to a 1.5-fold, p value <0.05) (Fig 2D, Fig S2B). On the other hand, 6 proteins exhibited induced or reduced abundance in the nucleoplasmic fractions upon rDNA damage (Fig 2D and Table S5). Notably, the nucleolar protein TCOF1 showed increased abundance in the nucleoplasmic fraction of GRASP cells, indicating an early stage of nDDR activation upon rDNA damage (Fig 2D). Thus, while we did not observe drastic dynamic changes in the nucleolar proteome, we did observe an initial trend in alterations of nucleolar-nucleoplasmic protein distribution at early stages of nDDR.

### DSBs at rDNA cause nucleolar replication stress

We next assessed the impact of rDNA damage on rDNA replication by examining markers of single stranded DNA (ssDNA) and replication stress: pRPA32 (S33) and (S4/S8), phosphorylated by ATR by DNA-PKs respectively, by performing co-IF with UBF. A specific induction of pRPA32 (S33) and not S4/S8 was observed in nucleoli after doxycycline treatment for 24h (Fig 3A-B). pRPA32 (S33) was recently shown to be detected when co-transcriptional R-loops at transcription termination sites (TTS) of highly active genes collide head-on (HO) with replication forks, while pRPA32 (S4/S8) marks co-directional (CD) transcription-replication conflicts (TRCs)^16^. The distinct staining pattern of pRPA32 (S33) at the nucleoli in GRASP cells indicate that rDNA DSBs lead to HO-TRCs at rDNA.

**Figure 3.**
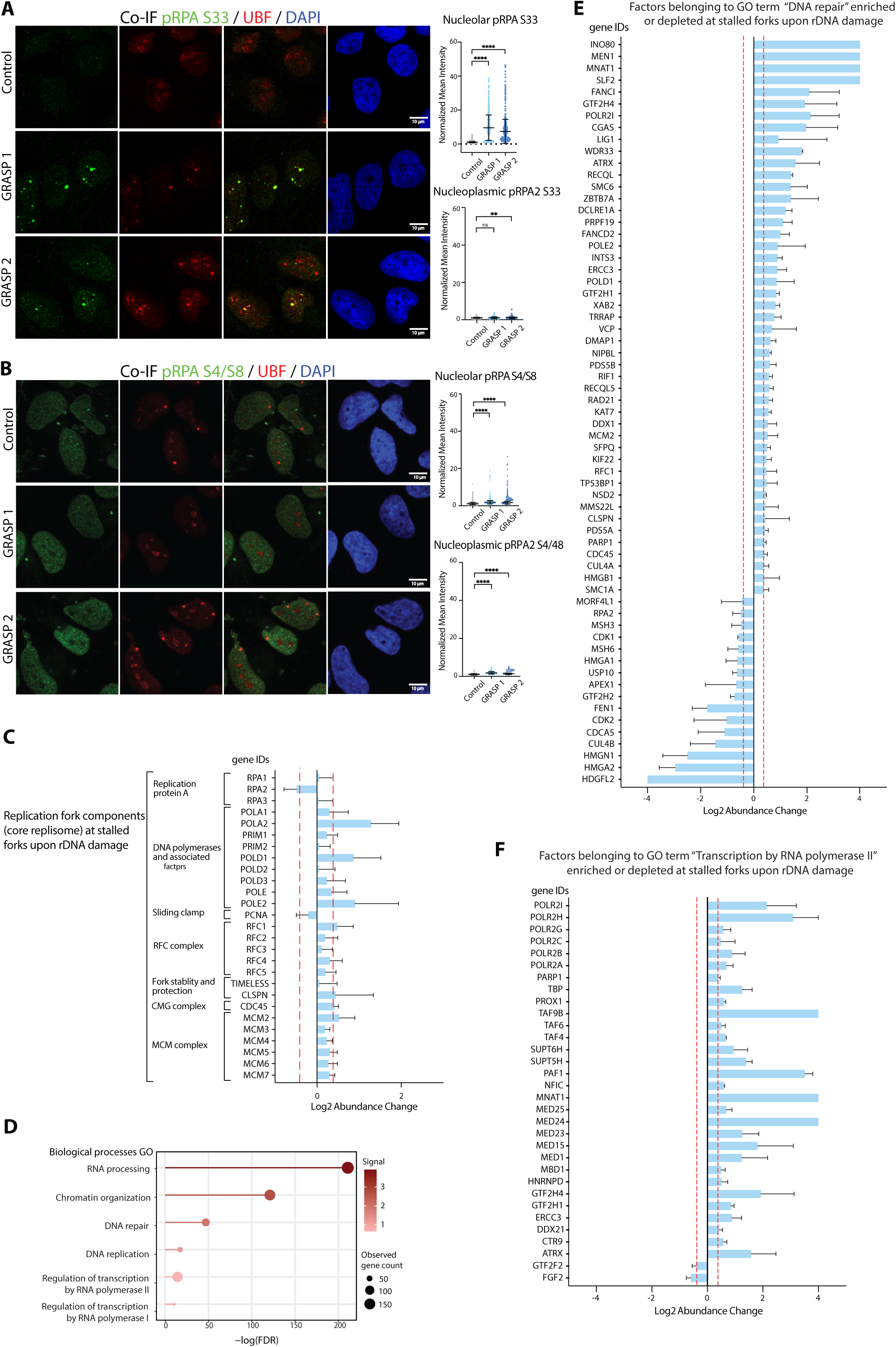
rDNA damage induces nucleolar replication stress. **A.** Co-IF analysis of pRPA32 (S33) and UBF in OVCAR8 control and GRASP cells treated with doxycycline (100 ng/ml) for 24h. Representative images from three independent experiments. Quantitation of signal intensity was performed using CellProfiler, normalized to the mean value of control cells per experiment. Over 400 cells were analyzed per condition across three independent experiments. Error bars represent mean ± SD. Statistical analysis was performed using unpaired t test, *ns* indicated a non-significant p-value greater than 0.05, ** p-value ≤ 0.01, **** p-value ≤ 0.0001. **B.** Co-IF analysis of pRPA32 (S4/8) and UBF as in (**A**). **C**. AniPOND-MS (accelerated native isolation of proteins on nascent DNA followed by MS analysis) to analyze the proteome at stalled replication forks upon rDNA damage. A schematic of EdU labelling of nascent DNA and aniPOND procedure is provided in Fig S3A. Log_2_ enrichment of core replisome proteins at stalled replication forks in GRASP cells treated with doxycycline (100 ng/ml) for 24h, compared with active replication forks in control cells. **D**. 358 and 188 proteins were identified to be commonly enriched or depleted, respectively (Table S6), at stalled replication forks in both GRASP clones with 1.3-fold change relative to EdU-labelled active forks in control cells, from two biological replicates per condition. Lollipop plot of the biological processes of proteins enriched at the stalled fork in GRASP cells. **E**. Log_2_ change of factors enriched/ depleted at stalled replication forks in GRASP cells compared with active replication forks in control cells, that belong to GO term “DNA repair” and **F**. GO term “Transcription by RNA Pol II”.

Since the majority of pRPA32 (S33) replication stress signal is observed at the nucleoli in GRASP cells (Fig 3A), we next performed aniPOND-MS (accelerated native isolation of proteins on nascent DNA followed by MS analysis) to analyze the proteome at stalled replication forks in the two GRASP cell lines, using two biological replicates. This method retrieves proteins associated with replication forks via capture of EdU-labelled DNA, which is then biotinylated and subjected to streptavidin retrieval (Fig S3A). The cells were incubated with doxycycline for 24h, then pulsed with EdU (10 μM) for 15 min with and without a subsequent thymidine (Thy 10 μM) chase for 1h to distinguish proteins that are enriched at EdU-labelled DNA fragments, including PCNA, by comparison to chromatin (Thy chase) (Fig S3B). In control cells, we identified 300 proteins as having been upregulated 1.3-fold or more in EdU-only (active forks) samples over Thy chase control (Table S6), of which 32 are known components of the replisome or are associated with DNA replication and DDR^17^. These factors included PCNA, the replication factor C (RFC) complex, and replicative polymerases delta 1 (POLD1) (Fig S3C). We also intersected our data with previously reported replisome datasets ^17–21^ that used various retrieval methods in various cell types and identified common components with all interested datasets that include factors involved in chromosome organization, cell cycle regulation and DNA repair (Fig S3D-F).

Alterations in the proteome at stalled replication forks in GRASP cells following rDNA damage were then compared to EdU-only samples of active forks in control cells. We proposed that this comparison enables the identification of replication stress response factors at the rDNA, since the induction of pRPA32 (S33) in GRASP cells was specific to the nucleoli (Fig 3A) and the cell cycle analysis indicated only a mild effect of the S-phase progression by 24h of doxycycline induction of rDNA DSBs (Fig 1I). We first interrogated the data for the core replisome, chromatin remodelling factors and DDR (Fig 3C) and identified that within 24h of rDNA damage, components of the core replisomes were still bound to replication forks.

In total, 358 and 188 proteins were identified to be commonly enriched or depleted, respectively (Table S6) at stalled replication forks in both GRASP clones with 1.3-fold change relative to EdU-labeled active forks in control cells. GO analysis of the chromatin composition at stalled forks in GRASP cells identified enriched signatures linked to chromatin organization, DNA replication and repair, RNA processing, Pol I and Pol II transcription (Fig 3D-F). Notably, an intersection analysis of the enriched proteins at stalled forks in GRASP cells with the nucleolar proteome of OVCAR8 cells identified a 68.1% (244/358) overlap (Table S7) suggesting that the local nucleolar proteome plays a critical role in responding to DSBs at rDNA. This localized response includes chromatin remodelling events, DDR, increased Pol II gene expression and RNA processing activity at rDNA (Fig 3E-F).

To validate the signatures observed in the aniPOND data, we performed a small-scale CRISPR-Cas9 knock out (KO) screen of 34 factors that belonged to enriched GO terms of chromatin remodeling, regulation of gene expression and RNA processing, to assess if their depletion affects nucleolar morphology as an indicator of defects in ribosome biogenesis and nucleolar function (Fig 4). OVCAR8 and an independent cell line OVCAR4 cells expressing Cas9 were transfected with sgRNAs (2-3 individual sgRNA per gene) for 96h. Cells were then fixed and co-IF staining for FBL and NPM was performed followed by counterstaining with DAPI and high throughput imaging. The data show that targeting factors enriched in our aniPOND screen caused alterations in nucleolar morphology, including fragmented FBL and/ or reduced and diffused NPM staining, indicating defects in ribosome biogenesis (Fig 4A-B). A significant concordance in the impact of gene KO on nucleolar morphology was observed between OVCAR8 and OVCAR4 cells (Fig 4C). Thus, our aniPOND screen uncovers factors involved in maintaining the delicate balance that coordinates Pol I transcription with DNA replication and repair at the rDNA loci and provides a valuable resource for investigating the molecular response to nucleolar replication stress.

**Figure 4.**
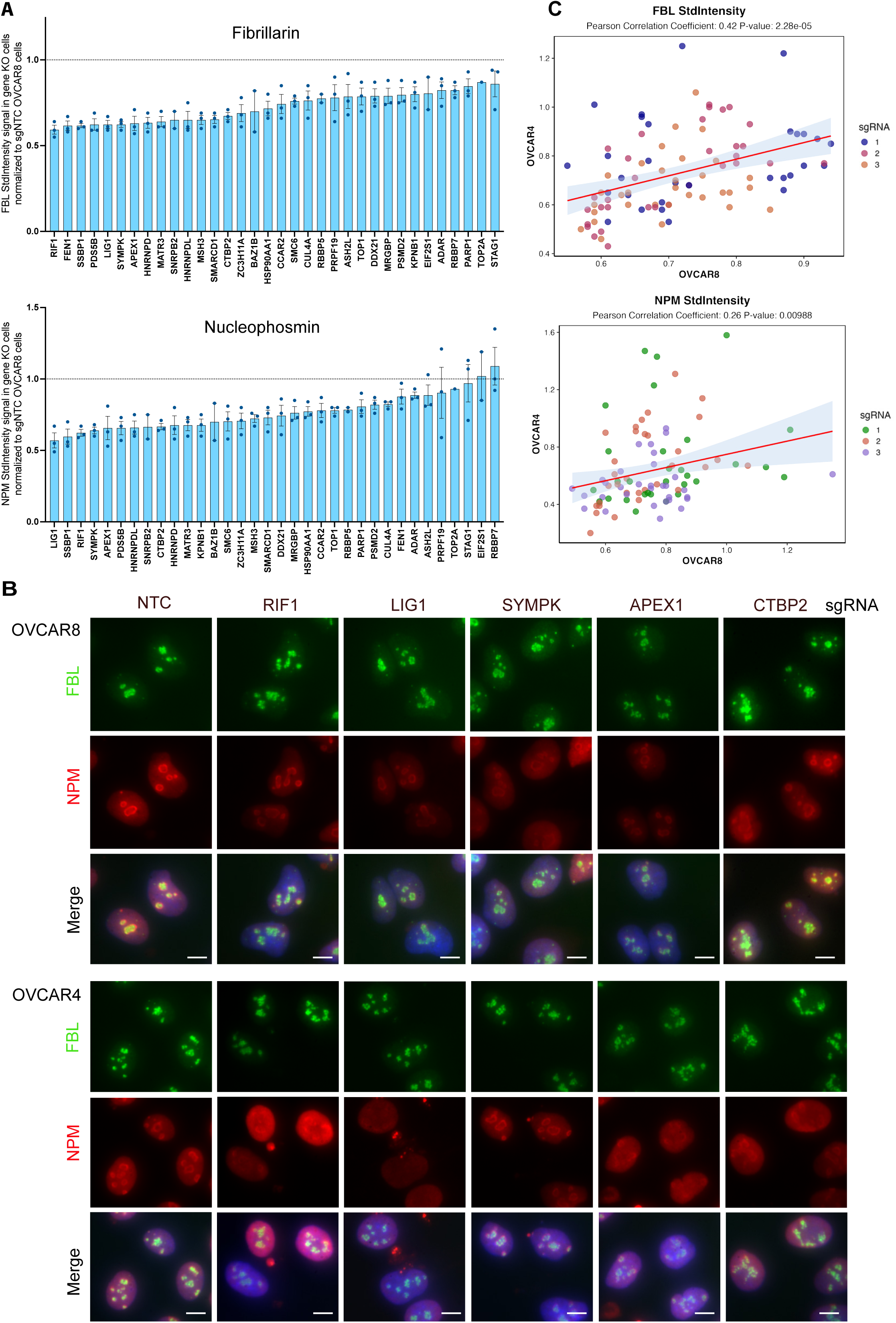
A boutique CRISPR-Cas9 screen of nucleolar stress factors OVCAR8 and OVCAR4 cells expressing Cas9 cell were plated in 384-well plates and reversed transfected with 3 individual sgRNAs per gene (Table S13) targeting 34 genes and non-targeting control (NTC) sgRNA, in 2 replicate wells per sgRNA. Following 96h, co-IF for FBL and NPM and counterstaining with DAPI was performed. High-through imaging was performed using a Thermo Fisher Cellomic CX7 LZR. For each well, 20 fields were captured across all channels using the 40X objective. sgRNAs that caused over 70% reduction in cell viability were excluded from the analysis. **A**. Images were analyzed using CellProfiler to measure the standard intensity (the standard deviation of pixel intensity) within regions marked by NPM and FBL. The median signal per field was taken, and the average across all fields per well was calculated and normalized to sgNTC. The normalized value of two technical wells was averaged and plotted with GraphPad Prism with each dot representing data from individual sgRNA. Error bars ± SEM. **B**. Representative co-IF images of FBL and NPM staining. Scale bar = 10 μm. **C**. A Pearson correlation coefficient of FBL and NPM StdIntensity measurements for FBL and NPM as in (A) for OVCAR8 and OVCAR4 cells.

### Increased Pol II transcriptional activity upon rDNA damages facilitates rDNA repair

Since R-loops are linked to HO-TRCs and signatures of Pol I and Pol II transcription and RNA processing activity are enriched at stalled replication forks upon rDNA damage (Fig 3D, 3F and Fig S3G), we performed co-IF for R-loops and UBF to assess the levels of nucleolar R-loops in GRASP cells. The nucleoli are naturally enriched with R-loops produced by Pol I and Pol II, which mediate strand-specific transcription of the IGS to produce sense intergenic non-coding RNAs (sincRNAs) and antisense intergenic ncRNAs (asincRNAs), respectively^22^. Pol II-transcribed asincRNAs form an antisense R-loop shield that limits the synthesis of Pol-I-dependent sincRNAs, which can compromise nucleolar condensates and abrogate nucleolar organisation^22^.

Treatments of OVCAR8 cells with the Pol I inhibitor Actinomycin D (Act D) and the Pol II inhibitor 5,6-Dichloro-1-β-D-ribofuranosylbenzimidazole (DRB) significantly reduced R-loops levels (Fig 5A-B). Interestingly, inhibiting Pol II activity significantly reduced nucleolar R-loops levels in GRASP cells by comparison to inhibiting Pol I transcription, indicating that the nucleolar R-loops species present upon rDNA damages are mainly produced by Pol II. In agreement with this, chromatin immunoprecipitation (ChIP) showed that the active form of Pol II pS2 was enriched at the IGS amplicon regions (Fig 5C-D). In contrast, Pol I localized primarily to the promoter and the 5’ external transcribed spacer (ETS) of the 47S rRNA coding region (Fig 5C). Upon the induction of rDNA DSBs, increased Pol II binding to IGS regions correlated with increased ratio of asincRNAs relative to sincRNAs at the IGS regions flanking the DSB in GRASP versus control cells (Fig 5E). On the other hand, decreased Pol I binding at the rDNA promoter region was observed, which corresponds to a decrease in Pol I transcription of the 47S pre-rRNA (Fig 1C). Thus, the results show that rDNA damage reduced Pol I transcription but increased Pol II transcription at IGS and facilitated R-loops formation.

**Fig 5.**
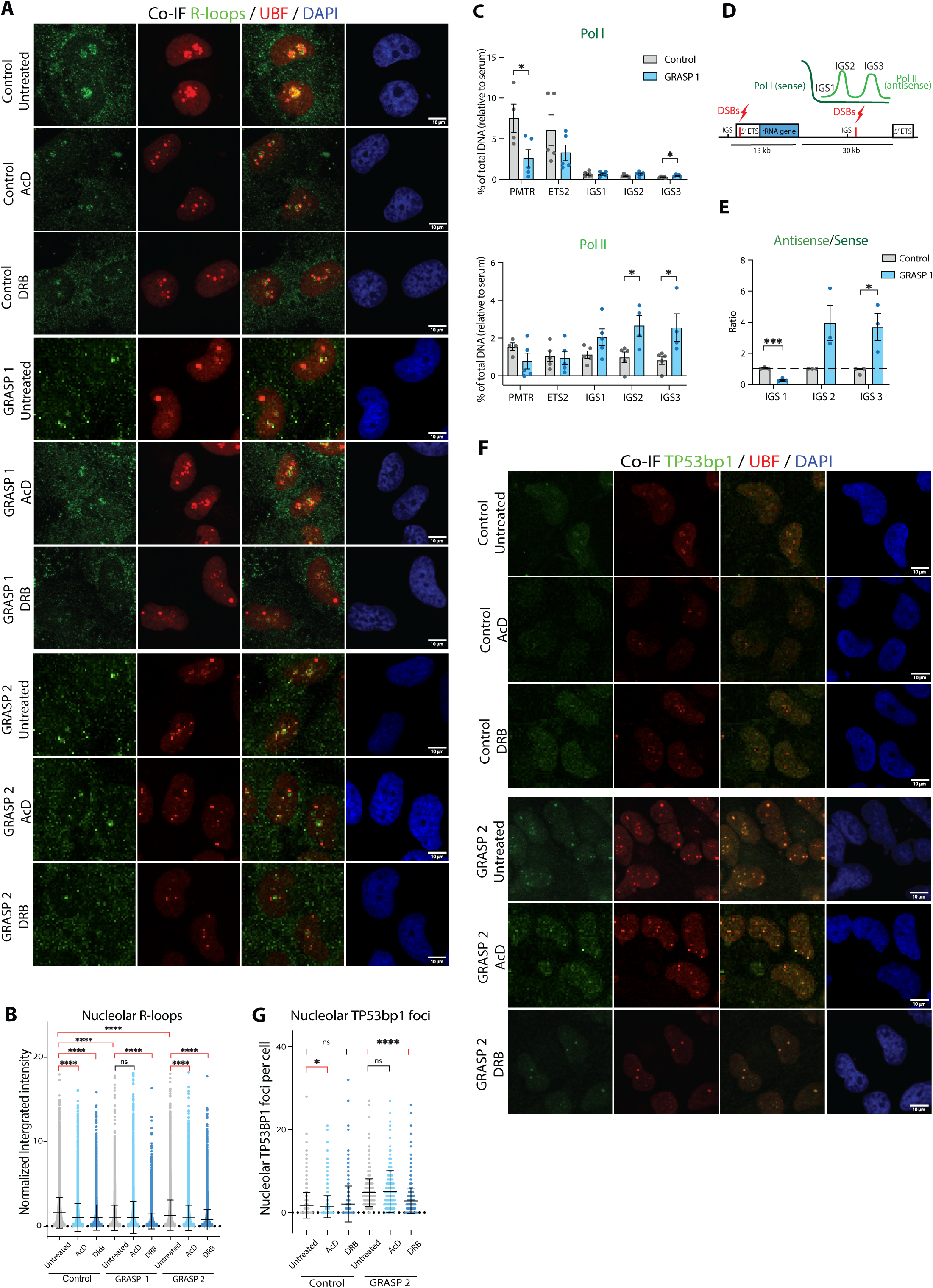
Pol II transcription-drives nucleolar R-loop formation and facilitates TP53BP1 recruitment upon rDNA damage. **A.** Co-IF analysis of R-loops and UBF in control and GRASP cells treated with doxycycline (100 ng/ml) for 24h and either vehicle, 10 nM Actinomycin or 100 µM DRB for the last 4 hours before harvest. Representative images of three biologically independent experiments. **B**. Quantitation of R-loops signal intensity was performed using CellProfiler. The integrated intensity was normalized to the corresponding median value of control cells per experiment. Error bars represent mean ± SD. Statistical analysis was performed using unpaired t test; *ns* indicated a non-significant p-value ≤ 0.05; **** is p-value ≤ 0.0001. RNAse H control condition for the specificity of S6.9 antibody in detecting R-loops is presented in Fig S4. **C**. ChIP-qPCR analysis of Pol I and Pol II binding to rDNA in GRASP and control cells treated with doxycycline (100 ng/ml) for 24h. Error bars represent mean ± SEM of *n*=5. Statistical analysis was performed using unpaired t test; * is p-value ≤ 0.05. **D**. A schematic of nucleolar Pol II at rDNA intergenic spacers (IGSs) producing antisense intergenic ncRNAs due to DSB at IGS. **E**. Strand-specific RT–qPCR (ss-RT) showing the levels of sense (Pol I) and antisense (Pol II) intergenic ncRNAs and at various IGS sites in GRASP and control cells treated with doxycycline (100 ng/ml) for 24h. Error bars represent mean ± SEM of *n*=3. Statistical analysis was performed using unpaired t test; * is p-value ≤ 0.05. **F**. Co-IF analysis of TP53BP1 and UBF in GRASP 2 cells treated as in (A). Representative images of three biologically independent experiments. **G**. Quantitation of TP53BP1 foci number was performed using CellProfiler. Error bars represent mean ± SD. Statistical analysis was performed using the unpaired t test; *ns* indicated a non-significant p-value greater than 0.05; * p-value ≤ 0.05; **** p-value ≤ 0.0001.

Since RNA transcribed by Pol II at DSB sites is known to play a crucial role in recruiting TP53BP1 to DSBs^23^, we investigated if inhibiting Pol II activity could affect TP53BP1 foci formation in GRASP cells. TP53BP1 foci were significantly decreased in both the number and intensity within the nucleolus specifically after Pol II inhibition with DRB treatment and not Act D treatment (Fig 5F-G). This suggests that nucleolar Pol II transcription upon rDNA damage drives R-loop formation to promote DNA repair and nDDR.

### RAD54L exhibits a synthetic interaction with Pol I transcription inhibition

To functionally assess interactions between DDR pathways and Pol I transcription, we undertook a synthetic lethal boutique CRISPR-Cas9 screen (Fig S5A and Fig 6A) targeting 15 DDR genes against Pol I inhibitors CX-5461^24^ and BMH-21^25^ and 9 other chemotherapeutic drugs, including classic Topoisomerase (TOP) I and II poisons, DDR inhibitors, the G4 stabiliser pyridostatin (PDS). These15 DDR genes spanning various DNA repair mechanisms such as HR, NHEJ, MMEJ (Microhomology-mediated End Joining), NER (Nucleotide Excision Repair), are pivotal protein mediators essential for DDR and genomic stability.

**Figure 6.**
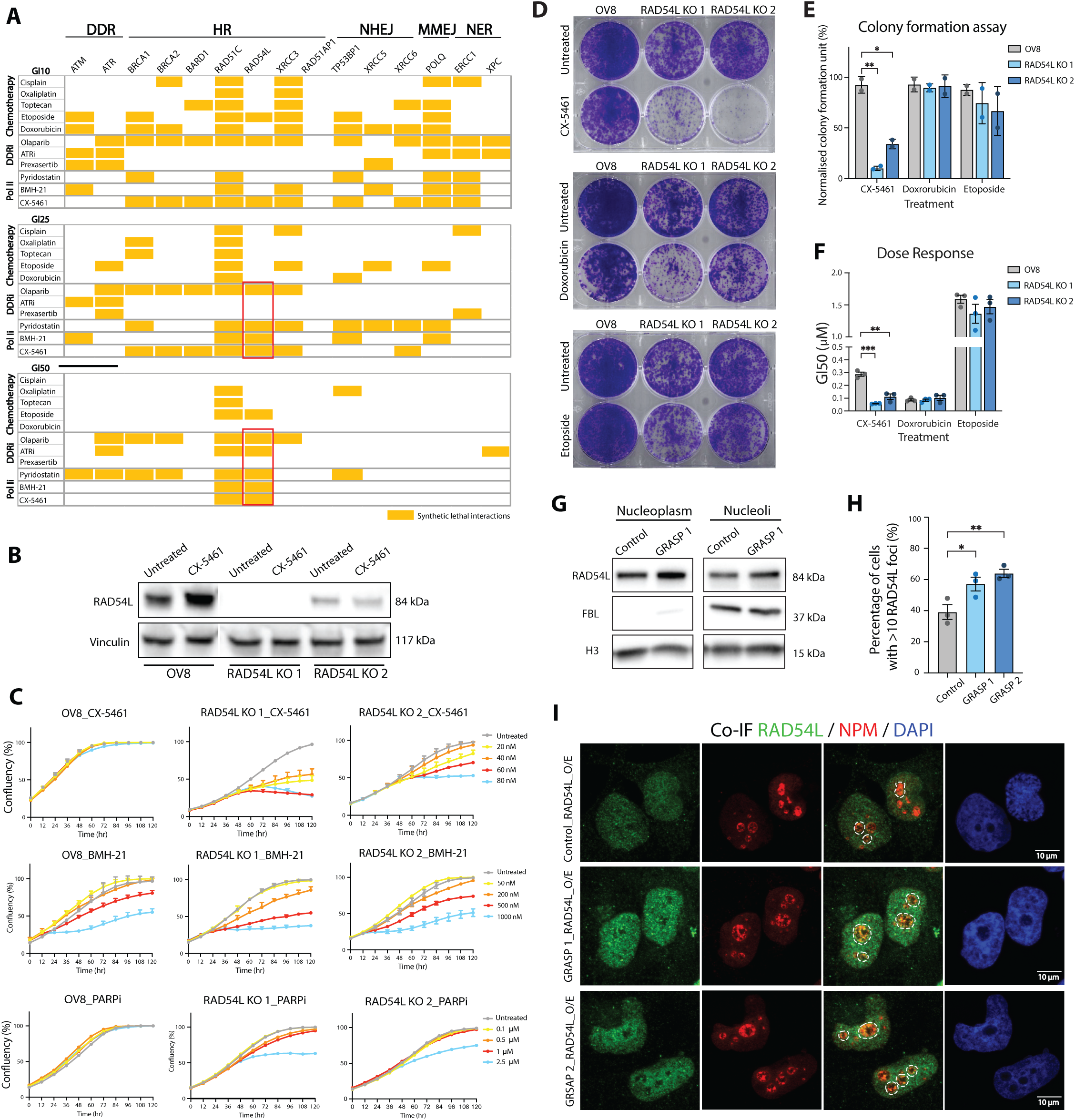
RAD54L exhibits synthetic lethal interactions with inhibitors of Pol I transcription. **A.** A snapshot of significant synthetic lethal (SL) interactions of DDR gene KO in OVCAR8 cells treated with 11 chemotherapeutics, DDR inhibitors, and Pol I inhibitors using three doses per drug for 72 hours (6 technical replicates per condition). SL interactions were assessed by calculating the absolute growth difference and the relative growth difference for each drug in gene KO relative to Cas9 control cells (Fig S5A). A stringent 15% threshold for absolute and 85% for relative differences in cell number was applied to identify significant SL interactions (highlighted in yellow). **B**. Western blot analysis of RAD54L expression in parental OVCAR8 cells and two derivative RAD54L KO cells with or without treatment with 1 μM CX-5461 for 3h. **C**. Cell confluence measured using IncuCyte live-cell imaging of OVCAR8 and RAD54L KO cells treated with different concentrations of CX-5461, BMH-21 and olaparib as indicated. Live-cell imaging was preformed every 12 hours for 5 days. Data shown is a representative experiment of three biological replicates. Error bars represent mean ± SEM of 3 technical replicates. **D-E**. Colony formation assay in parental OVCAR8 and RAD54L KO cells treated with 20 nM CX-5461, 10 nM Doxorubicin, or 200nM Etoposide for five days. Following another 7 days of incubation with drug-free median, cells were fixed and stained with 0.1% crystal violet solution. Representative images of wells are shown. % Occupancy per well was measured using Image J software and normalized to corresponding untreated controls. Error bars represent mean ± SEM of *n*=3. Statistical analysis was performed using an unpaired student t-test. * p-value ≤ 0.05, ** p-value ≤ 0.01. **F**. Sensitivity of RAD54L KO cells to CX-5461 and Top II poisons Etoposide and Doxorubicin were compared to parental OVCAR8 cells. Cells were treated with increasing doses for 72h then fixed and stained with DAPI to acquire the cell number. Growth inhibition dose response curves were plotted and GI50 doses were generated using GraphPad Prism 10. Error bars represent mean ± SEM of *n*=3. Statistical analysis was performed using an unpaired student t-test; ** p-value ≤ 0.01, *** p-value ≤ 0.001. **G**. Western blot analysis of RAD54L in nucleolar and nucleoplasmic fractions of control OVCAR8 cells and GRASP 1 cells treated with 100 ng/ml doxycycline for 24 hours. FBL and Histone H3 was used as a loading control within each fraction. **H**-**I** Co-IF analysis of RAD54L and NPM in RAD54L-overexpressing OVCAR8 and GRASP cells treated with doxycycline (100 ng/ml) for 24h. Representative images from three independent experiments. Cells with more than 10 RAD54L foci in the nucleolus (marked by NPM) were scored as RAD54L foci-positive cells. At least 100 cells per condition were scored, and the percentage of nucleolar RAD54L foci-positive cells are shown. Error bars represent mean ± SEM of 3 independent experiments. Statistical analysis was performed using an unpaired student t-test. * p-value ≤ 0.05; ** p-value ≤ 0.01.

Briefly, OVCAR8-Cas9 cells transfected with crRNAs for 96h, then replated into 96 well plates and either treated with DMSO or with pre-determined GI10, GI25 and GI50 doses of each drugs for 72h (Table S8). Cells were then fixed, stained with DAPI and imaged to determine cell viability. To characterise synthetic lethal (SL) interactions of drugs with DDR gene KO, we calculated the absolute growth difference and the relative growth difference for each drug in gene KO cells relative to Cas9 control cells (Fig S5A). A stringent 15% threshold for absolute and 85% for relative differences in cell viability was applied to identify significant SL interactions. A snapshot of SL interactions of gene KO with drug was generated (Fig 6A). As expected, we observed that *BRCA1/2* KO confer sensitivity to olaparib and chemotherapeutics. CX-5461 is a dual inhibitor of RNA Pol I and TOP II activities^24,26–28^, however, by comparison to other TOP2 poisons like doxorubicin and etoposide, CX-5461 acts as a DNA structure-driven TOP2-poison at transcribed regions bearing G4 structures with preferential activity at rDNA loci^26,29^. In this screen, CX-5461 exhibited a unique profile compared to other drugs showing SL interactions with HR and NHEJ factors and PolQ at the low GI10 10nM dose. The HR repair signature becomes more dominant with increasing doses of CX-5461. RAD54L emerged as the target demonstrating the strongest, specific SL interactions with CX-5461 and BMH-21. Notably, RAD54L also enhanced the sensitivity to olaparib and PDS. Thus, the data suggest that RAD54L may play an important role in mediating response to DNA damage at rDNA or G4-enriched regions.

To confirm the SL interaction of RAD54L and inhibitor of Pol I transcription, we established RAD54L KO OVCAR8 cell lines (Fig 6B). Proliferation and clonogenic assays confirmed that loss of RAD54L led to increased sensitivity towards Pol I inhibitors CX-5461 and BMH-21 and to PARPi olaparib (Fig 6C-F) compared to parental OVCAR8 control cells. However, RAD54L KO cells remained equally sensitive to TOP II poisons as control cells (Fig 5D-F). This unique sensitivity to Pol I inhibitors suggests that RAD54L may play a role in nDDR or replication stress.

Interestingly, western blot analysis of subnuclear fractions of OVCAR8 control and GRASP cells showed that RAD54L resides in the nucleoli and the nucleoplasm under normal condition and its levels were increased in both fractions in GRAPS cells upon rDNA damage (Fig 6G). Similarly, RAD54L foci increased in the nucleoli and the nucleoplasm of OVCAR4 cells treated with CX-5461 for 3h (Fig S5B). Moreover, control and GRASP cells overexpressing RAD54L exhibited an increase in % of cells with RAD54L foci at the nucleoli upon rDNA damage relative to control cells (Fig 6H-I), supporting a role for RAD54L in mediating nDDR.

### RAD54L play distinct roles in signalling DDR, restraining replication forks and mediating HR DNA repair

RAD54L is a DNA-dependent ATPase that regulates RAD51 during HR repair at the different stages of filament stabilization, displacement loop (D-loop) formation, and removal from paired strands^30^. The RAD54 family of proteins are members of the Swi2/Snf2 family of motor proteins that utilize energy from ATP hydrolysis to translocate on dsDNA. RAD54 and RAD54L stabilize the interaction of RAD51 with ssDNA tracts. RAD51 then carries out a search for homologous dsDNA sequences and promotes invasion of target duplex leading to the exchange of DNA strands that forms heteroduplex DNA within an intermediate called the D-loop. Subsequent stages of the recombination process result in repair of damage without loss or rearrangement of DNA sequences. RAD54L then removes RAD51 from dsDNA thus eliminating unnecessary recombination and counteracting the genome-destabilizing effects of direct binding of RAD51 to dsDNA.

To assess the DDR to CX-5461 in the absence of RAD54L, western blotting for DDR proteins was performed (Fig 7A). As previously shown, CX-5461 treatment of OVCAR8 cells resulted in activation of DDR signalling including phosphorylation of ATM, ATR, CHK1, CHK2 and RPA32 (S4/S8). However, the loss of RAD54L significantly reduced DDR activation compared to parental OVCAR8 cells. Notably, RPA32 phosphorylation is increased in RAD54L KO cells treated with CX-5461, suggesting that the loss of RAD54L leads to increased levels of ssDNA lesions and replication stress following CX-5461 treatment and that it is an essential component for DDR signalling.

**Figure 7.**
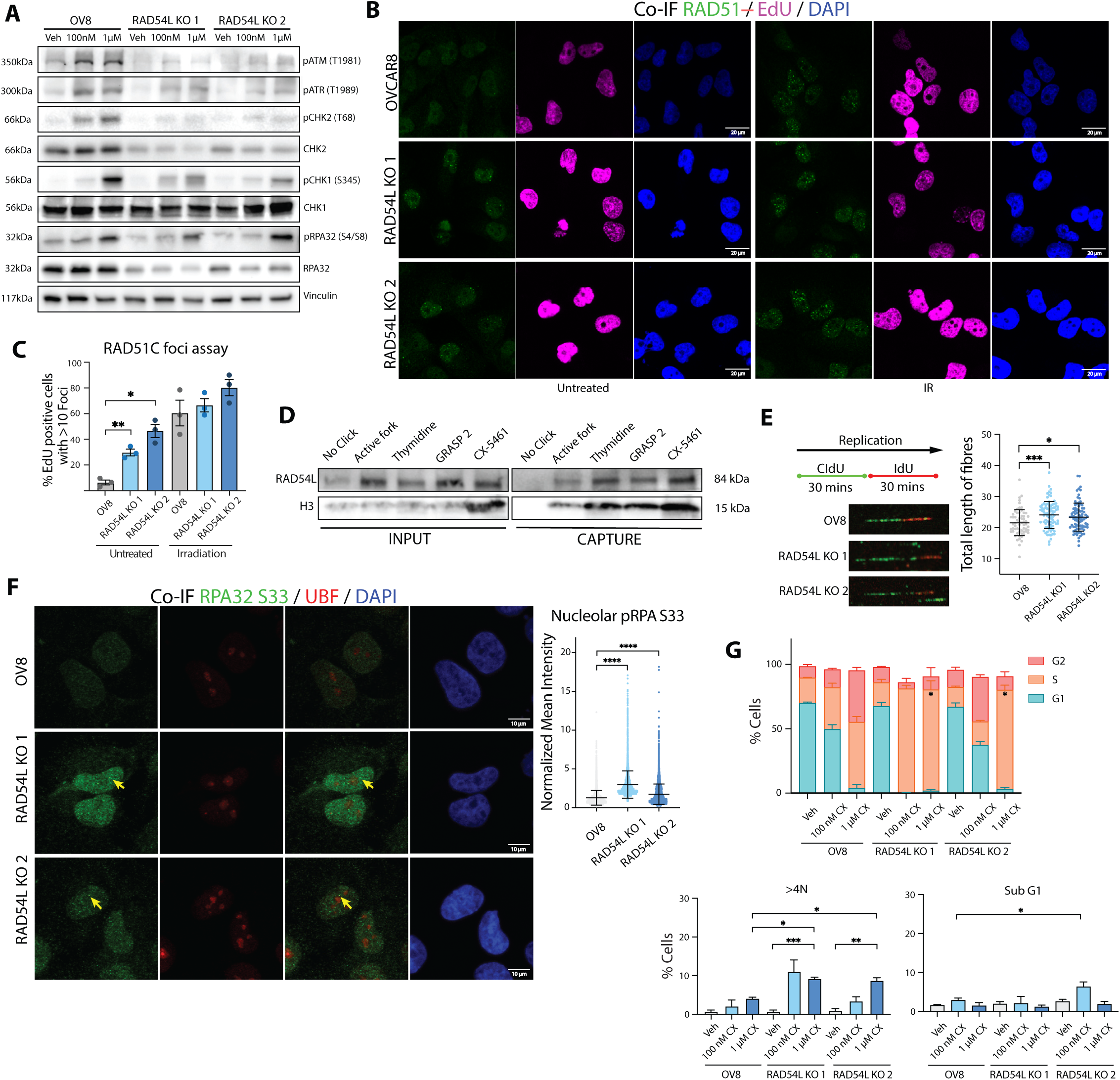
RAD54L plays distinct roles in signalling DDR, restraining replication forks and mediating HR DNA repair. **A**. Western blot analysis of key DDR proteins in parental OVCAR8 and RAD54L KO cells treated with CX-5461 for 24h as indicated. Blots are representative of three independent biological replicates. **B**. IF analysis of RAD51 in parental OVCAR8 and RAD54L KO cells, either untreated or following ionising irradiation (IR, 10 Gy for 20 minutes). Cells were labelled with 10 μM EdU prior to IR to mark S-phase cells, allowed to recover for 6 hours, then fixed and stained. **C**. S-phase cells (EdU-positive) with more than 10 RAD51 foci were marked as RAD51 foci-positive cells. Manual scoring was performed with at least 100 cells per condition. Error bars represent the mean ±SEM of three biological replicates. Statistical analysis was performed using an unpaired t-test; * ≤ 0.05, ** ≤ 0.01. **D**. Western blot analysis of aniPOND capture showing enrichment of RAD54L in control and GRASP 2 cells treated with doxycycline (100 ng/ml) for 24h and OVCAR8 cells treated with 1 µM CX-5461 for 3h. **E**. Schematic of labelling for DNA replication assessment using fibre assays. Cells were labelled with 10 μM CldU for 30 minutes followed by 10 μM IdU for 30 minutes, then harvested. Representative fibres from parental OVCAR8 and RAD54L KO cells are shown. Track lengths were measured using ImageJ. Quantitation includes at least 75 fibres per biological replicate, with a total of at least 200 fibres analyzed across three replicates. Statistical analysis was performed using an unpaired student t-test. * p-value ≤ 0.05, *** p-value ≤ 0.001. **F**. Co-IF analysis of pRPA32 (S33) and UBF in parental OVCAR8 and RAD54L KO cells. Representative images from three independent experiments. Quantitation of signal intensity of pRPA/UBF colocalized regions was performed using CellProfiler and normalized to median value of OVCAR8 cells per experiment. Over 400 cells were analyzed per condition across three independent experiments. Error bars represent mean ± SD. Statistical analysis was performed using an unpaired student t-test; **** p-value ≤ 0.0001. **G**. Cell cycle analysis of BrdU and PI labelling in parental OVCAR8 and RAD54L KO cells following 72h of CX-5461 treatment (100 nM or 1 µM). Histogram plots show % cell populations in G1, G2M and S-phase, SubG1 and cells with >4N DNA content. Error bars represent mean ± SEM of *n*=3. Statistical analysis was performed using an unpaired t-test, * p-value ≤ 0.05, ** p-value ≤ 0.01, *** p-value ≤ 0.001.

In agreement with RAD54L’s role in displacing RAD51 filaments, RAD54L KO cells exhibit higher levels of RAD51 foci at baseline levels (Fig 7B-C), suggesting an increase in RAD51 binding to DNA resulting from an imbalance between RAD51 and RAD54L translocase activity. Interestingly, RAD54L depletion does not affect RAD51 binding to damaged DNA in ionizing radiation (IR)-treated cells (Fig 7B-C). This suggest that RAD54L loss does not affect RAD51 binding to DNA damage sites but the increase in RAD51 foci at baseline levels indicate that RAD54L limits RAD51 binding at undamaged DNA. The data highlights an important role for RAD54L in removing RAD51 from dsDNA, which is important for preventing HR in situations/locations where such recombination is unnecessary and potentially detrimental, such as the rDNA loci.

In addition to its role in HR DNA repair, RAD54L was more recently shown to play a role in restraining replication fork progression and promoting fork reversal to maintain fork stability during replication stress^31^. Indeed, RAD54L binding to replication forks was enhanced following treatment with CX-5461, and RAD54L KO cells exhibited faster rates of replication fork progression (Fig 7D-E). Moreover, loss of RAD54L led to increases in RPA32 (S33) levels, including foci formation at the nucleoli and nucleoplasm indicating increased rDNA and global replication stress (Fig 7F).

While the loss of RAD54L did not affect cell cycle profiles at baseline conditions, RAD54L KO cells treated with CX-5461 exhibited S-phase delay and increased proportions of cells with > 4N content, suggesting failed cytokinesis (Fig 7G). Thus, in the absence of RAD54L unresolved CX-5461-mediated replication stress leads to prolonged S-phase, and failed cytokinesis upon entering mitosis with damaged DNA.

### RAD54L secures fidelity of the rDNA loci

We next examined the impact of RAD54L loss on nucleolar morphology and function by performing co-IF analysis of FBL and NPM (Fig 8A-B) and qRT-PCR for Pol I transcription activity (Fig 8C). The data revealed enlarged nucleoli with increased FBL fragmentation and a significant reduction in 47S rRNA levels in RAD54L KO cells compared to control cells. Further investigation showed elevated levels of R-loops in RAD54L KO cells (Fig 8D-E), suggesting disruptions in transcription and replication at rDNA loci. Additionally, the loss of RAD54L led to a pronounced increase in γH2AX levels, indicative of enhanced DNA damage both within the nucleolus and globally (Fig 8F-G). These findings identify an unrecognized role for RAD54L in the nDDR surveillance pathway and underscore its critical role in maintaining nucleolar integrity and genome stability.

**Figure 8.**
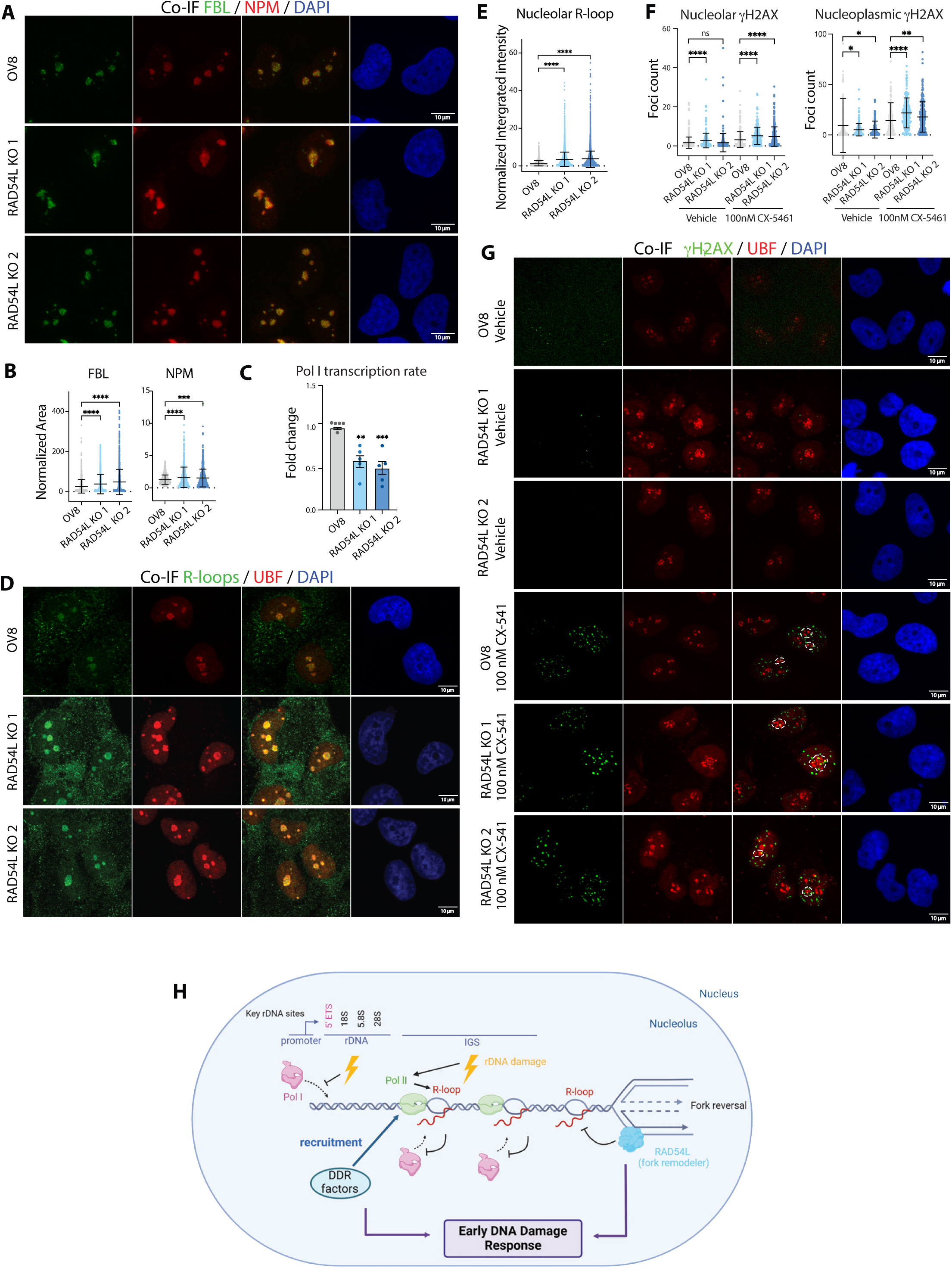
Loss of RAD54L leads to nucleolar stress. **A**. Co-IF analysis of FBL and NPM in exponentially growing parental OVCAR8 and RAD54L KO cells. Representative images from three independent experiments are shown. **B**. Quantitation of FBL and NPM area was performed using CellProfiler, normalized to the median value of OVCAR8 cells per experiment. Over 400 cells were analyzed per condition across three independent experiments. Data are shown as mean ± SD. Statistical analysis used an unpaired t-test; **** p-value ≤ 0.0001. **C**. Pol I transcription levels were assessed by quantitative real-time PCR (qRT-PCR) analysis using primers specific to the 5’ETS of the 47S rRNA precursor. Expression levels were normalized to Vimentin mRNA and shown as fold changes relative to control cells. Data represent means from three independent biological replicates; error bars indicate mean ± SEM. Statistical analysis was performed with an unpaired t-test. ** p-value ≤ 0.01, *** p-value ≤ 0.001. **D**. Co-IF analysis of R-loops and UBF in parental OVCAR8 and RAD54L KO cells. Representative images of three independent biological experiments are shown. **E**. Quantitation of R-loops signal intensity was performed using CellProfiler, with integrated intensity normalized to the median value of OVCAR8 cells per experiment. Error bars represent mean ± SD. Statistical analysis used an unpaired t-test; **** is p-value ≤ 0.0001. **F-G**. Co-IF analysis of γH2AX and UBF in OVCAR8 and RAD54L KO cells. Representative images from three independent biological experiments are shown. Quantification of nucleolar and nucleoplasmic γH2AX foci count was performed using CellProfiler. Data are presented as mean ± SD. Statistical analysis was conducted using unpaired t-test; * p-value ≤ 0.05, ** p-value ≤ 0.01, **** p-value ≤ 0.0001. **H**. Schematic illustrates the process by which nucleolar Pol II transcription drives the formation of nucleolar R-loops, facilitating the recruitment of DDR factors and triggering the early stages of the DNA damage response (DDR). RAD54L also plays a protective role at rDNA restraining replication forks, sensing replication stress and activating DDR.

Altogether, our data highlights an important protective role for RAD54L at resolving replication stress such as TRCs at highly transcribed and G4 and R-loops-enriched regions including the rDNA loci (Fig 8H). Importantly, our data of characterizing the proteome of stalled replication sites upon rDNA damage provide a platform for investigating TRCs, which were recently identified as a source of synthetic lethality with PARPi^32^. As such identifying mechanisms that enhance TRCs can provide therapeutics opportunities.

## Discussion

DNA repair and replication of the rDNA repeats must be orchestrated with ongoing transcription. Our findings showed an enrichment of factors involved in DNA damage and repair, chromatin assembly, transcription (particularly Pol II), and RNA processing at replication forks following the induction of rDNA damage. Notably, around 68.1% of the factors detected at stalled replication forks are part of the nucleolar proteome. The data therefore suggest that local nucleolar surveillance networks play a crucial role in the immediate or early response to rDNA damage. We further identified a new role for nucleolar Pol II in producing R-loops that promote TP53BP1 recruitment for rDNA repair. The data highlights a role for Pol II in early response to rDNA DSBs (Fig 8H).

Here, we define a nucleolar surveillance network of proteins that responds to rDNA damage at rDNA loci. This network is likely to play a critical role in ensuring rDNA genome stability and provide a valuable tool for understanding the response to nucleolar replication stress. This is important considering recent data showing replication stress as the underlying mechanism of synthetic lethality with PARPi and a therapeutic vulnerability in cancer cells^32^. Indeed, we identify RAD54L as a new component of nDDR that is important for resolving R-loops at rDNA and protecting rDNA stability, in addition to its known roles in HR repair and restraining replication forks.

Characterizing nDDR provides broad implications for health and disease. Mutations in genes involved in ribosome biogenesis and assembly account for the pathogenesis of a heterogeneous group of diseases called ribosomopathies^33^. Mutation of *TCOF1* is the primary mechanism for Treacle Collins Syndrome (TCS), a well-known ribosomopathy^33,34^. The critical role of TCOF1 in rDNA repair and ribosome biogenesis has been considered the primary cause of the extensive apoptosis of neuroepithelial cells associated with the pathogenesis of TCS^35^. The nDDR and nucleolar stress response are also increasingly recognized as contributors to neurodegenerative diseases^36^. Moreover, the nDDR pathway is targeted by infectious viruses that interact with TCOF1 to inhibit its function, thereby inhibiting nDDR and silencing ribosome biogenesis^37^. Many viruses target the nucleolus to interact with nucleolar proteins to facilitate replication, transcription, and assembly of viral particles^38^. Thus, further exploration of the critical factors involved in nDDR could provide valuable insights into the molecular mechanisms driving ribosomopathies, viral infections and age-related diseases, such as neurodegenerative disorders.

Oncogene-driven cancer cells have higher demands for Pol I transcription and ribosome synthesis compared to normal cells, making them more sensitive to rDNA instability, nDDR activation and nucleolar stress^3^. Therefore, targeting early nDDR factors could selectively impair cancer cell growth, offering a promising anti-cancer therapeutic strategy. Several chemotherapy drugs, such as cisplatin and TOP II poisons, can induce the nucleolar stress response and initiate nDDR by inducing rDNA damage^33,39^. However, these drugs also cause global DNA damage, which causes toxicity and severe side effects. As a dual inhibitor of RNA Pol I and TOP II activities^24,26–28^, CX-5461 induces nucleolar stress and nDDR^24,29,40^. CX-5461 is synthetic lethal with HR and other DNA repair pathways^40,41^. A recent Canadian Phase Ib clinical trial^42^ of CX-5461 (Pidnarulex) in heavily pre-treated solid tumor patients, including known *BRCA1/2* germline mutation showed that out of 15 patients who were evaluable for drug response, 40% achieved clinical benefit, with stable disease (SD) being the best therapeutic response observed. Among the patients with SD were 5 ovarian cancer patients with mutations in HR genes, who had previously failed platinum chemotherapy and PARPi treatments, with 2 patients maintaining SD for at least 6 months following treatment with CX-5461^42^, highlighting the potential of harnessing nDDR in overcoming resistance to traditional treatments. Overall, this study opens new avenues for developing nDDR-targeting therapies and deepens our understanding of the molecular mechanisms underlying nDDR.

## Methods

### Cell lines and culture conditions

The human HGSOC OVCAR8 cells expressing inducible CRISPR-associated protein 9 (Cas9) were transduced with two single guide RNAs (sgRNA1 and sgRNA2) (Table S9) using the pCLIP-dual-SFFV-Puromycin lentiviral system. Clonal cell lines were then derived and referred to as “GRASP”. crRNAs targeting RAD54L (Table S13) were used to KO RAD54L in OVCAR8 cells and two clones were selected for this study. OVCAR4, OVCAR8 Cas9, OVCAR8 control (expressing pCLIP-dual-SFFV-Puromycin empty vector), GRASP cell lines and OVCAR8 RAD54L KO cell lines were grown in RPMI-1640 medium supplemented with HEPES, 10% (v/v) foetal bovine serum (FBS), 2 mM GlutaMAX (Gibco, 35050061), cultured at 37 °C in a humidified incubator with 5% CO_2_ and maintained in culture for a maximum of 8–10 weeks. The identity and individuality of cell lines were routinely confirmed by a polymerase chainreaction (PCR) based short tandem repeat (STR) analysis using six STR loci. Mycoplasma testing was routinely performed by PCR.

### Compounds

CX-5461 was purchased from MedChemExpress and prepared as a10 mM stock in 50 mM NaH_2_PO_4_. VE-821 (ATRi), BMH-21, prexasertib, pyridostatin and olaparib were purchased from Selleckchem, with 10mM stock were prepared in DMSO. Etoposide was also obtained from Selleckchem and prepared at 100mM in DMSO. Topotecan was acquired from Selleckchem, with a 1 mM stock were prepared in 0.9% saline. Pharmacy-purchased cisplatin, oxaliplatin and doxorubicin were dissolved in 0.9% saline.

### Cell proliferation assays

Dose-Response curves for assessing drug sensitivity were generated by plating cells in 96-well plates for 24h, then treating them with either vehicle or increasing concentrations of drugs for 72h. Cells were then stained with DAPI (Thermo Fisher, #D1306) and scanned using IncuCyte S3 ZOOM imaging system (Essen BioScience) to determine cell counts. GI doses were generated using GraphPad Prism 10. The dose of drug that inhibits cell growth by 10%, 25%, 50%, 75% and 90% (GI10, GI25, GI50 and GI90) at 72h was identified (Table S8). For real-time proliferation assays, cells were seeded in 96-well plates, incubated overnight and then treated with drugs for up to 7 days. Confluency was measured every 12 hours using the IncuCyte S3 ZOOM imaging system to assess cell growth dynamics.

### Cell cycle analysis

For cell cycle analysis using 5-bromo-2 –deoxyuridine (BrdU) incorporation, cells were pulse labelled with 10 μM BrdU (Sigma-Aldrich, B5002) for 30 minutes then washed twice with PBS, collected, fixed in 80% ice-cold ethanol. Fixed cells were stored at 4 °C until further processing. For BrdU staining, fixed cells were pelleted and incubated in 1 mL of 2N HCl containing 0.5% (v/v) Triton X-100 for 30 minutes. Cells were then pelleted and washed in 1 mL of 0.1 M Na_2_B_4_O_7_.10H_2_O, pH 8.5 (Sigma-Aldrich, #S9640-500G) and sequentially incubated for 30 minutes with anti-BrdU (Abcam, #ab6326) and FITC-conjugated anti-mouse IgG (Jackson ImmunoResearch, #515-095-003) diluted in PBS containing 2% FBS and 0.5% Tween-20. Cells were then washed with PBS containing 2% FBS and incubated in 10 µg/mL propidium iodide (PI) solution at room temperature for 15 minutes. Flow cytometric analysis was performed using the LSRFortessa™ Cell Analyzer (BD Biosciences), and cell cycle profiles was analysed with Flowlogic software (Version 10, Inivai Technologies).

### Clonogenic assays

Cells were seeded in 6-well plates for 24 hours and following treatment with drugs for 5 days, the media was aspirated, and cells were washed with PBS. Drug-free media was then added and cells were incubated for additional 7 days, then fixed with 100% methanol for 1 h, stained with 0.1% (w/v) crystal violet solution (Sigma-Aldrich, C6158) for 1 h, rinsed with H_2_O and air-dried. Colonies were imaged and quantitated using ImageJ software.

### Western blotting

Cell lysates were separated by SDS-PAGE and transferred onto Immobilon-P poly-vinylidene fluoride (PVDF) membranes (MERCK Millipore, IPVH00010) via electrophoresis. Proteins were detected using Enhanced Chemiluminescence (ECL) detection (Bio-Rad, #1705062). Antibodies details are listed in Table S11.

### Label-free proteomics and Mass spectrometry

Cells were lysed with guanidium chloride lysis buffer containing 6 M guanidinium chloride, 100 mM Tris pH 8.5, 10 mM tris(2-carboxyethyl)phosphine (TCEP), and 40 mM 2-chloroacetamide. Samples were boiled at 95°C for 10 min and sheared with 26G needles. For proteomics, 100 µg of protein sample was precipitated with acetone and methanol at –20 °C overnight and then resuspended in 50 mM Tris-HCl pH 8 after washing twice with 80% (v/v) acetone. 20 μg of each protein sample were digested by Trypsin/Lys-C Mix, Mass Spec Grade (0.4 g/uL) (Promega, V5071) at 37 °C, 1,000 rpm overnight. The digestion was terminated the following day by adding Trifluoroacetic acid (TFA) (Sigma-Aldrich, #302031) to a final concentration of 1% (v/v). Samples were desalted using of C18 Empore™ SPE Disks (Merck, #66883-U), and eluted with 60% (v/v) acetonitrile and 5% (v/v) formic acid. Organic solvents were removed by speed vacuum evaporation, then samples were resuspended in 15 μL of 1% (v/v) formic acid and subject to LC-MS analysis.

For phosphoproteomics studies, 300 μg of proteins were precipitated, washed, and digested by Trypsin/Lys-C Mix as described above. The phosphorylated peptides were enriched using the PureCube Fe(III)-nitrilotriacetate MagBeads (Cube Biotech, #31501-Fe) by rotating at room temperature for 1h, and the beads were washed with 80% (v/v) acetonitrile and 0.1% (v/v) TFA, and phosphopeptides were eluted with 50% (v/v) ACN and 2.5% (v/v) ammonium hydroxide (Sigma 338818) through home-made C8 stage tips.

Samples were analyzed using the Thermo Scientific™ UltiMate™ 3000 RSLCnano system coupled with an Orbitrap Fusion™ Lumos™ Tribrid™ Mass Spectrometer in Proteomics facility at St Vincent’s Institute. Peptide mixtures were loaded onto a Thermo Scientific™ Double nanoViper PepMap™ Neo column (75 µm x 50 cm). Peptides were identified from the mass spectrometry spectra output using Proteome Discoverer 3.1 or MaxQuant 2.4.2.0 software with Perseus 1.6.15.0. For phosphoproteomics analysis, peptides with phosphate localization probability higher than 0.75 were used for further analysis. The significantly differentially expressed proteins or phosphorylated peptides were identified by Students t tests, defined by p value ≤0.05 and absolute fold change ≥1.5.

### RNA extraction and Reverse transcription and qPCR

RNA was extracted using a NucleoSpin RNA Mini kit for RNA purification (Macherey-Nagel, #740955.50). First-strand cDNA synthesized was performed using random hexamer (Promega, C1181) and Superscript IV reverse transcriptase (Invitrogen, #18080044). The reactions were incubated at 25 °C for 10 min, 37 °C for 60 min, and 70 °C for 15 min. Regarding strand-specific qPCR analysis, each cDNA sample was amplified using primers directed at the strand-specific transcript of interest as described in^22^. This included ss_Tag and hIGS_forward primers for sense transcripts, ss_Tag and hIGS_reverse primers for antisense transcripts, and 7SK forward primers as a control for 7SK sense transcripts (Table S10).

Quantitative PCR (qPCR) was performed using FAST SYBR Green dye (Applied Biosystems, #4385610) on the StepOnePlus real-time PCR system (Applied Bio-systems). The qPCR reactions were carried out with an initial denaturation at 95 °C for 5 minutes, followed by 39 cycles of 95 °C for 5 seconds and 60 °C for 30 seconds. Primer sequences are detailed in Table S11.

### Immunofluorescence

For IF assays combined with EdU labelling, cells were first incubated with 10 μM EdU for 30 minutes to label S-phase cells prior to drug treatment. Cells were fixed in 4% paraformaldehyde (PFA) for 10 minutes at room temperature, permeabilized with ice-cold 0.3% Triton X-100 in PBS for 10 minutes, and washed with PBS. Blocking was performed with 5% goat serum and 0.3% Triton X-100 in PBS for 30 minutes at room temperature. Subsequently, cells were incubated with primary and secondary antibodies (Table S12) at 37 °C for 1h in a humidified chamber. EdU incorporation was visualized using a Click-IT reaction buffer containing 100 mM Tris, pH 8.5, 10 nM Alexa Fluor 647-azide (Invitrogen, #A10277), 1 mM CuSO4, and 100 mM ascorbic acid. This reaction was carried out at room temperature for 30 minutes, followed by PBS washes. Nuclei were counterstained with Vectashield mounting media containing DAPI. For IF staining using the S9.6 (R-loops) antibody, cells post-PFA fixation were permeabilized with 100% methanol for 10 minutes and 100% acetone for 1 minute on ice. For RNAse H control condition, after PBS washing, RNase H treatment was performed by applying a final concentration of 50 units /mL to each slide and incubating at 37°C in a dark, humid chamber for 3h (Fig S4). Images were acquired using a Leica STELLARIS 5 Cryo Confocal Light Microscope equipped with 63x magnification with a water-based objective, Z-step set to 1. Data analysis was conducted using with ImageJ and CellProfiler.

### AniPOND

AniPOND was performed based on previously published methods^43,44^ with modifications. After 100 ng/ml doxycycline treatment for 24h, GRASPs and control cell were incubated with 200 µM EdU for 15 minutes. Control cells were also subjected to a thymidine chase by replacing the EdU-containing media with fresh media containing 10 μM thymidine (Sigma-Aldrich, #T1895) for 1h. After EdU labelling or Thymidine chase, the media was aspirated. Ice-cold nuclei extraction buffer (20mM HEPES pH 7.2, 50mM NaCl, 300 mM Sucrose, and 3 mM MgCl_2_ in H2O with 0.5% IGEPAL CA630) was added, and cells were then scraped. The lysates were incubated on ice for 15 minutes, nuclei were collected by centrifugation at 500 g for 10 minutes at 4 °C. The nuclei pellets were then resuspended in 10 mL of ice-cold click reaction mixture (25 μM biotin-azide (Jena Bioscience, #CLK-1265-25),10mM (+)-sodium L-ascorbate (Sigma Aldrich, #A7631-25G), and 2 mM CuSO_4_ in PBS), and rotated at 4 °C for 1h, followed by centrifugation at 500 g for 10 minutes at 4°C. For “No Click” negative control, an equivalent volume of DMSO was used instead of biotin-azide. Following rotation, the Click-IT reaction mixture was centrifugated at 500 g for 10 minutes at 4 °C, and the pellets were resuspended in 0.5 mL of ice-cold buffer B1 (25 mM NaCl, 2 mM EDTA, and 50 mM Tris-HCl pH 8.0 in H_2_O with 1% IGEPAL CA630). Samples were rotated at 4°C for another 30 minutes and centrifugation at 500 g for 10 minutes at 4°C. This process was repeated twice.

The samples were then sonicated using a sonicator (Bioruptor™, #UCD-200) at low power for 35 cycles (30 sec on and 30 sec off). The sonicated material was centrifuged at 15000g for 10 minutes at 4 °C, and the supernatant was collected. This process was repeated once, and the final supernatant volume was adjusted to 1 mL with ice-cold buffer B1. A 20 µL aliquot was saved as the INPUT for western blot analysis. The remaining samples were incubated overnight at 4°C with streptavidin-coated beads (Thermo Fisher, #20353). Beads were washed in ice-cold buffer B1 five times. The washed beads were then incubated at 100 °C for 15 minutes in a 1:1 volume 2×Laemmli sample buffer (Bio-RAD, #1610737). Finally, supernatants as the CAPTURE samples were collected after a brief spin, stored at –20 °C, and used for either Western blotting or MS preparation.

For MS analysis, in-gel digestion was performed. CAPTURE samples were loaded in SDS-PAGE gels and electrophoresed until the protein entered the top 0.5 cm of the gel. The gel region was excised, and the gel was stained with 0.5% SimplyBlue™ SafeStain coomassie blue G-250 (Invitrogen™, #LC6065) in 10% acetic acid for 5 minutes. Excess stain was removed by rinsing briefly with MilliQ water, and the gel was destained with 40% HPLC-grade methanol/ 10% acetic acid for 10–20 minutes until faint bands were visible. Subsequently, the gel was destained overnight in MilliQ water.

The protein bands were further excised into small pieces (1-2 mm pieces) using a clean razor blade and transferred to microcentrifuge tubes. The gel was destained with 800 µL of destaining buffer (50 mM TEAB, 100% ACN, mixed 1:1) three times, the first destain lasted 3h, followed by two additional washes of 30 minutes each. After removing the destaining buffer, the gel pieces were dehydrated with 800 µL of 100% ACN for 30 minutes. The gel pieces were reduced by adding 100 µL of 10 mM TCEP in 50 mM TEAB and incubating for 45 minutes at 55 °C. After removing the excess solution, 100 µL of 55 mM iodoacetamide in 50 mM TEAB was added for alkylation in the dark for 30 minutes at room temperature. The gel pieces were washed three times with 500 µL of 50 mM TEAB, then dehydrated with 800 µL of 100% ACN for 15 minutes, and air-dried for 10 minutes. Proteins were digested overnight Trypsin LysC Mix at 37 °C with shaking at 800-1000 rpm. The resulting supernatant was collected and prepared for MS analysis as described above. All the buffers in this experiment included protease inhibitors (Focus Bioscience, # HY-K0011).

### Subnuclear Fractionation

Protocol was adapted from^45^. Cells were detached using with StemPro™ Accutase™ Cell Dissociation Reagent (Thermo Fisher, #A11105-01) at 37 °C for 10 minutes, and resuspended in culture medium, and washing twice with PBS. Cells were pelleted and resuspended in 5 mL Buffer A (20 mM Tris pH 7.4, 10 mM KCl, 3 mM MgCl_2_, 0.1% IGEPAL CA-630 and 10% glycerol) and lysed by sequential passage through a 21-gauge syringe three times and a 27-gauge syringe once. The lysate was centrifuged at 200 g for 5 minutes at 4 °C, and the pellet was resuspended in 3 mL Buffer B1 (0.25 M Sucrose, 10 mM MgCl_2_). This was overlaid onto 3mL Buffer B2 (0.35 M Sucrose, 0.5 mM MgCl_2_), and centrifuged at 1500 g for 5 minutes at 4°C. The nuclear pellet was resuspended in 3 mL Buffer C1 (0.5 M Sucrose, 3 mM MgCl_2_) and sonicated using a Branson™ Sonifier (3 round of 50% duty cycle, 10s on and 10s off). The sonicated material was layered onto Buffer C2 (1 M Sucrose, 3 mM MgCl_2_) and centrifuged at 2800g for 5 minutes at 4°C. The supernatant containing the nucleoplasm fraction was precipitated using acetone, while the nucleolar fraction was washed by resuspension in 0.5 mL Buffer C2 and centrifuging at 2800 g for 5 minutes at 4°C. The nucleolar pellets were analysed by western blot analysis or resuspended in 100 μL lysis buffer for MS analysis. All the buffers in this experiment included protease inhibitors.

### Chromatin Immunoprecipitation (ChIP)

After treatment with 100 ng/ml doxycycline for 24 hours, GRASP and control cells were fixed with 0.6 % PFA in medium at 37 °C for 10 minutes. The reaction was quenched by adding 0.125 M glycine at room temperature for 10 minutes. Cells were scraped with PBS, after washing twice, and then centrifuged at 3000g for 5 minutes. The cell pellets were resuspended in (10 mM Tris pH 7.4, 10 mM NaCl, 10 mM MgCl_2_, 0.5% IGEPAL CA-630 in DEPC-treated water, supplemented with protease inhibitor) and incubated on ice for 10 minutes. The lysate was centrifuged at 9000 g for 3 minutes at 4 °C, and the supernatant was discarded. The pellet was resuspended in MNase buffer and subjected to MNase digestion (1 gel units/μL, New England Biolabs, #M0247S) at 37 °C for exactly 10 minutes, with vortexing every 2.5 minutes. The digestion was terminated by adding 0.05 M EDTA, followed by incubation on ice for 5 minutes. The sample was centrifuged at 9000 g for 5 minutes at 4 °C. The pellet was resuspended with ChIP dilution buffer (167 mM NaCl, 1.67 mM Tris pH 8.1, 0.01% SDS, 1.1% Triton X-100 and 1 mM DTT in DEPC-treated water, supplemented with protease inhibitor) and sonicated using a Branson™ Sonifier (40% duty cycle, 3 rounds of 6s on and 30s off). The sonicated materials were centrifuged at 9000 g for 10 minutes at 4 °C, and supernatant was collected.

For the immunoprecipitation, DNA (100 µg) from each sample was diluted to 1mL with ChIP dilution buffer, and 6% of the sample was set aside as the INPUT control and stored at –80°C. 3-5 ug of Pol I and Pol II antibodies (Table S12) were used for each sample and incubated overnight at 4 °C. The antibody-lysate mixtures were captured with A/G magnetic beads (Thermo Fisher, #88803) and rotated for 2 hours at 4 °C. After incubation, the beads were washed with low salt buffer (150 mM NaCl, 20mM Tris pH 8.1, 2mM EDTA, 1% Triton X-100 and 0.1% SDS), high salt buffer (500 mM NaCl, 20mM Tris pH 8.1, 2mM EDTA, 1% Triton X-100 and 0.1% SDS), LiCl wash buffer (0.25 M LiCl, 10 mM Tris pH 8.1, 1mM EDTA, 1% sodium deoxycholate and 1% NP-40) and lastly twice with TE wash buffer (10 mM Tris pH 8.1, 1 mM EDTA). The chromatin was eluted from the beads by rotating the sample twice with elution buffer (100 mM NaHCO_3_ and 1% SDS in DEPC-treated water) for 15 min at room temperature. Input samples were thawed on ice and brought up to the same final volume as the samples with elution buffer. All samples were reverse cross-linked by incubation at 65 °C overnight with 200 mM NaCl.

DNA cleanup was performed by a 2h incubation at 45 °C after adding 9 mM EDTA pH 8.1, 36.4 mM Tris-HCl pH6.5 and 20 µg proteinase K. Samples were then mixed 1:1 with a combination of phenol, chloroform, isoamylalcohol (25:24:1) and centrifuged at 15000 g for 20 minutes at 4 °C. The upper aqueous phase (∼450 µL) was carefully collected, and 500 µL of isopropanol, 50 µL 3M sodium acetate and 1 µL of glycogen (20 mg/ml) were added. The mixture was centrifuged at 15000 g for 20 minutes at 4 °C, and the supernatant was discarded. The pellet was washed once with 70% ethanol and resuspend in 100 µL DEPC-treated water. qPCR analysis was conducted using primers listed in Table S12.

### DDR CRISPR-Cas9 synthetic lethal screen

For examining synthetic lethality of drugs with DDR genes, OVCAR8 cells expressing Cas9 were reversed transfected in using Dharmafect 4 (Horizon, #31985070) with crRNA duplexes (Millenium Science) (Table S13) and TracRNA (Horizon, #U-002000) or with sgRNA non-targeting control. After 24h, the transfection medium was replaced to growth medium, and cells were incubated for additional 72 hours to allow sufficient time for gene editing. Cells were then replated in 384-well plates using Biotek3 Liquid Handler and treated with 11 drugs at GI10, GI25 and GI50 doses in 3 replicates. Drug doses were determined from cell viability dose-response assays (Table S8).

Cells were then fixed, stained with DAPI and imaged using Cellomics CX7 LZR (Thermo Fisher) at x10 magnification. Cell survival was assessed based on cell number determined from DAPI-stained cell counts. Significant synthetic lethal (SL) interactions were assessed by firstly calculating the fold change in cell proliferation in drug-treated KO cells relative to DMSO-treated gene KO cells. Two factors were then calculated: SL_minus and SL_divid, for each gene KO and drug treatment. SL_divid is the percentage change of gene KO after drug treatment compared to the change in parental OVCAR8 Cas9 with the same treatment. To avoid identifying gene KO with a strong effect on cell proliferation alone, the absolute difference was also calculated between gene KO and OVCAR9 Cas9 after drug treatment (SL_minus). Therefore, the synthetic lethal hits were selected as >0.15 of SL-minus and <0.85 of SL_divid.

### Boutique CRISPR/Cas9 screen of nucleolar stress factors

OVCAR8 and OVCAR4 cell lines expressing Cas9 were plated in PhenoPlate™ 384-well microplates (Corning) and reversed transfected using Dharmafect 4 with 2-3 individual sgRNAs per gene and sgRNA non-targeting control (NTC) (Millenium Science), delivered using JANUS liquid handling robotic (Perkin Elmer) in duplicate plates. The plates were incubated at room temperature for 20 minutes. After 24h, 50 µL of fresh medium was added using Biotek3 Liquid handler, and plates were incubated at 37 °C in 5% CO₂ in a humidified incubator for additional 72h, followed by high throughput IF staining for NPM and FBL, which was performed as described above with appropriate volumes for 384-well plates. The primary antibodies for NPM and FBL were used at 1:2500 dilution for 90 minutes. The corresponding secondary antibodies at 1:1500 dilution were also incubated for 90 minutes (Table S11). A post-fixation step using 4% PFA was carried out followed by DAPI staining. High-through imaging was performed using a Thermo Fisher Cellomics CX7 LZR. For each well, 20 fields were captured across all channels using the 40X objective. Images were analyzed using CellProfiler, the median signal per field was taken, and the average across all fields per well was calculated and normalized to sgNTC. The normalised value of two technical wells was averaged and analyzed with GraphPad prism and R. Out of focus images and data for sgRNAs that caused over 70% reduction in cell viability were excluded from the analysis.

### DNA fibre analysis

Exponentially growing cells were pulse-labelled for 30 minutes with 50 μM CldU (Sigma-Aldrich, #C6891), washed three times with warm PBS and then incubated with 250 μM IdU (Sigma-Aldrich, 17125) for 30 min. Cells were the washed again in warm PBS, trypsinized and resuspended in ice-cold PBS at a concentration of 8 × 10^5^ cells/mL and spun down. The cell pellets were first resuspended in Buffer 1 (1:1 PBS:0.5% Trypsin without phenol red, Gibco, # 15400054) and heated at 50 °C for 10 seconds. Cells were then mixed with Buffer 2 (1.2% low melting agarose in PBS, Bio-Rad, #1613111) and immediately dispensed into DNA plug moulds (Bio-Rad, #1703713). The plugs were stored at 4 °C for at least 1 hour. Cell plugs were incubated in 250 μL of Buffer 3 (0.5 M EDTA pH8, 1% Sarkosyl and 2 mg/mL Proteinase K) overnight at 50 °C. After incubation, plugs were washed three times for 1h in 1 mL diluted Buffer 4 (1M Tris + 0.1M EDTA) and once for 3.5h. Following washing, Buffer 7 (0.5 M MES hydrate, pH5.5) was added, and the plugs were incubated at 68 °C for 20 minutes and then 42 °C for 10 minutes. Subsequently,1.5 μL of beta-agarase (New England Bio labs # M0392S) was added to the plug, which were then incubated overnight at 42 °C.

After incubation, sample were gently poured into the reservoir (Genomic Vision, RES-001). Fibres were combed using Fiber Comb combing machine (Genomic Vision) onto Combicoverslips (Genomic Vision, COV-002). The coverslips were glued onto Superfrost slides using non-fluorescent superglue. The slides were then dehydrated by baking at 65 °C for 2h and then denatured in 5 mL of freshly made 0.5 M NaOH + 1M NaCl for 8 minutes. The slides were washed with and then dehydrated by sequential ethanol incubations (70%, 90%, and 100%) for 5 minutes each, followed by air-drying. IF staining was performed as described above using primary antibodies: Rat anti-CldU and Mouse anti-BrdU, and appropriate secondary antibodies (Table S12). Imaging was performed using Leica Thunder Microscope with 40X Objective and fibre length was measured using ImageJ manually.

### Statistics and reproducibility

No statistical method was used to predetermine the sample size, but we routinely employed at least three biological repeats for each experiment, in each case scoring as many technical replicates as possible (typically several hundred cells). Randomization was not appropriate because samples were derived from specific genetic cell lines or treatment condition. The investigators were not blinded to allocation during the experiments and outcome assessment because all numerical data were software automated and independent of investigator subjectivity. All data presented in the manuscript are represented as the mean [plus or minus] SEM or SD and were analyzed using GraphPad Prism (version 10). The statistical test employed for each dataset has been specified in the figure legends.

## Acknowledgement

The authors would like to acknowledge Diannita Kwang, Dingyi Yu, Henry Beetham, the Biological Optical Microscopy Platform (University of Melbourne), Flow cytometry facility (SVI) and Mass Spectrometry facility (SVI) for technical support.

## Funding

This research was funded by Cancer Australia (2023/PCRS/0251), the Australian National Health and Medical Research Council (NHMRC Ideas grant #2029800) and Tour de Cure Foundation research grants. E.S. received support from the Victorian Cancer Agency (Research Fellowship MCRF19007). R.L. received support from SVI Rising Star Buxtons Fellowship.

J.K. received support from the 5Point Foundation (Christine Martin Fellowship).

The Victorian Centre for Functional Genomics (K.J.S.) is funded by the Australian Cancer Research Foundation (ACRF), Phenomics Australia, through funding from the Australian Government’s National Collaborative Research Infrastructure Strategy (NCRIS) program, the Peter MacCallum Cancer Centre Foundation and the University of Melbourne Collaborative Research Infrastructure Program.

---

**Figure S1.**
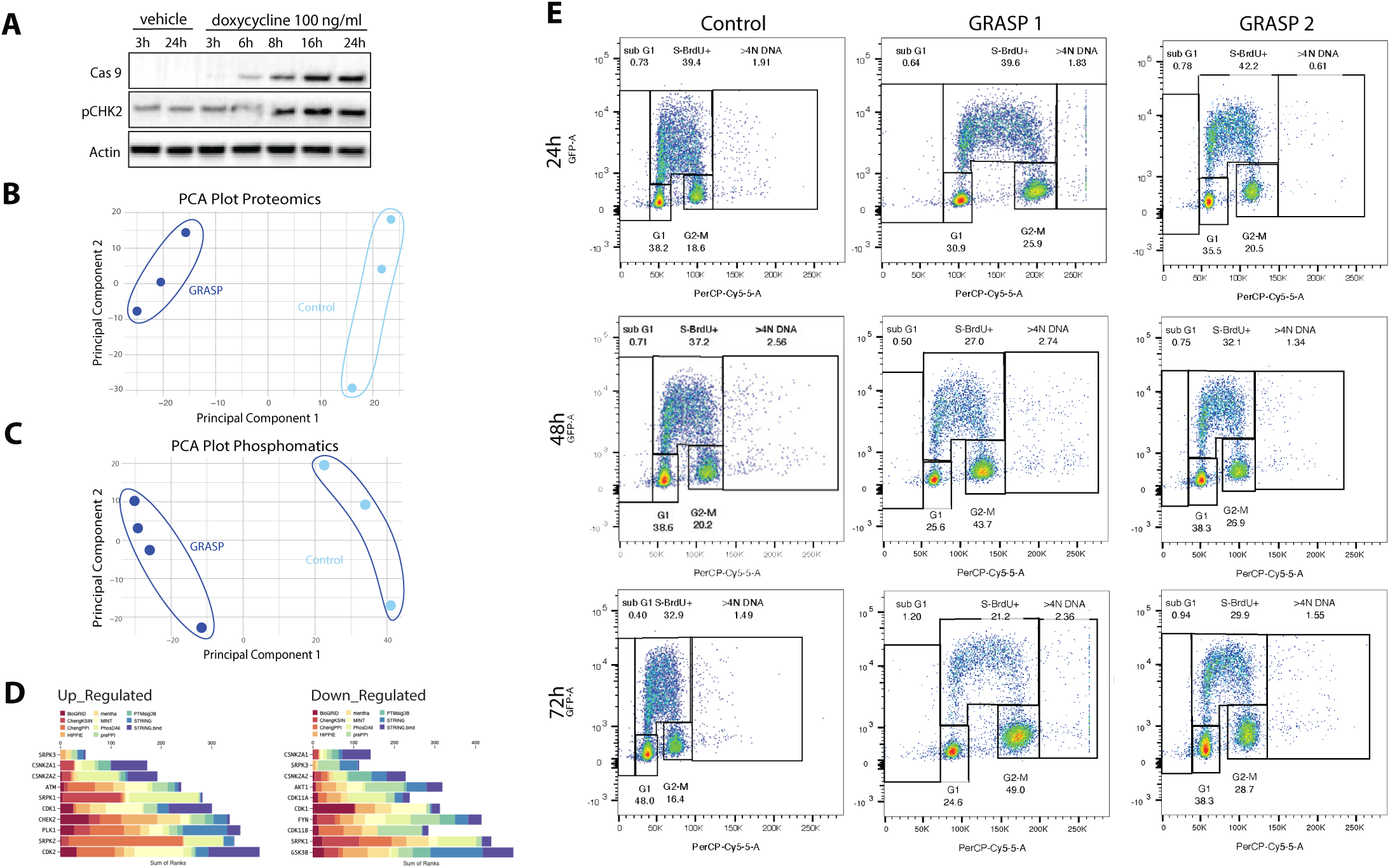
rDNA damage induces nucleolar DDR and G2/M cell cycle arrest. **A.** Western blot analysis of Cas9 protein and pCHK2 T68 expression in GRASP and control OVCAR8 cells treated with doxycycline over a time course as indicated. **B**. Principal Component Analysis (PCA) plot of proteomic and **C**. phosphoproteomics data of 4 replicate samples of GRASP 1 and control cells treated with 100 ng/ml doxycycline for 24h. **D**. Kinase enrichment analysis (KEA3) of the altered phosphoproteomic sites in GRASP cells compares control cells (Table S2) using MeanRank visualization from KEA3, indicating up-regulated and down-regulated activities of kinases. **E**. Analytical cell cycle analysis of BrdU incorporation as a function of DNA content using FACS. Cells were treated with doxycycline (100 ng/ml) for 24h, 48h and 72h and labelled with BrdU for 30 minutes prior to harvest. The boxes represent S-phase (BrdU-labelled), G0/G1, G2/M and Sub G0/G, along with cells with > 4N DNA content. The gating strategy for quantitating the percentage of cell populations is shown in the top panel. Representative data from three independent experiments (*n*=3).

**Figure S2.**
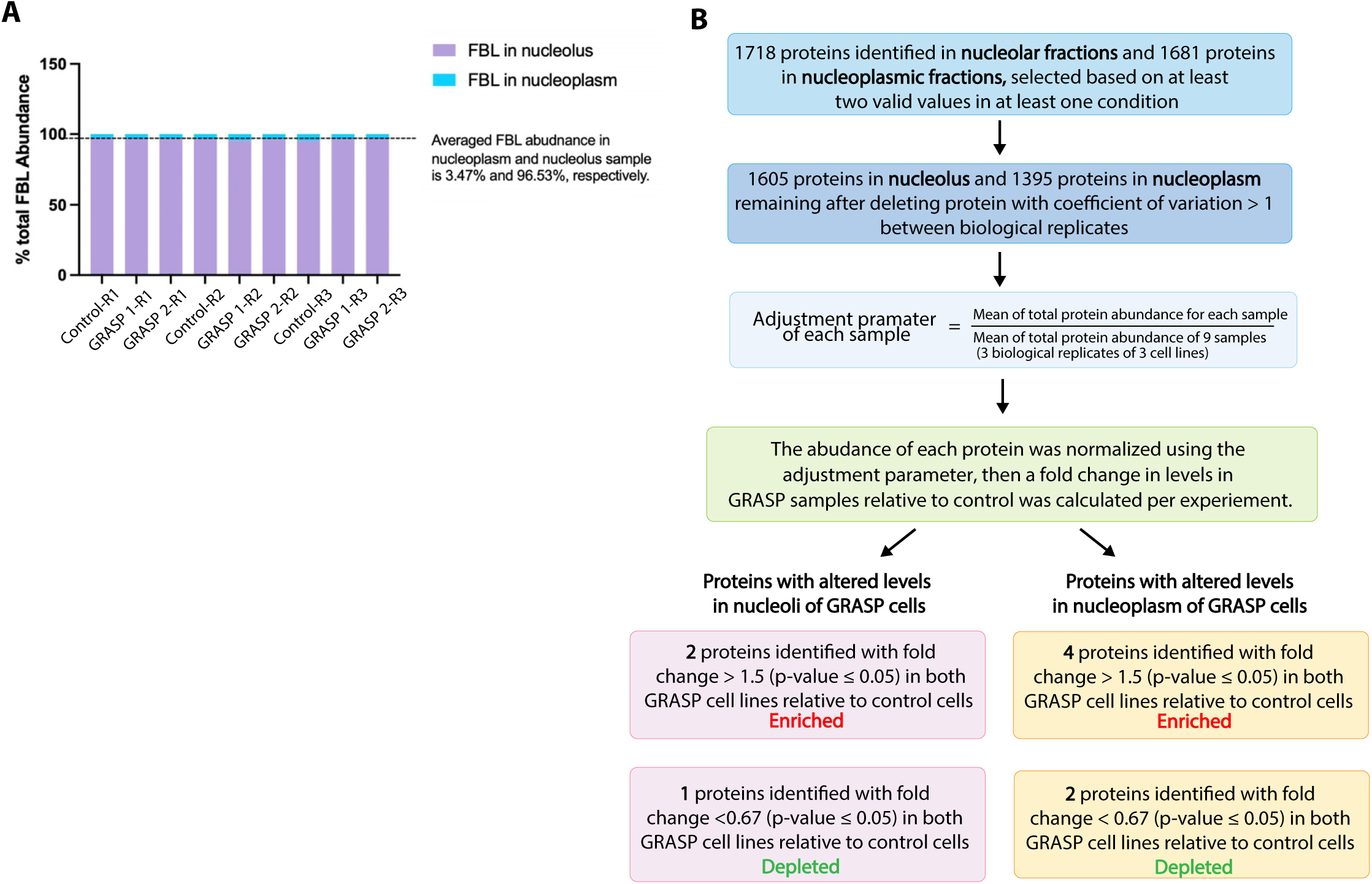
Subnuclear fractionation-proteomic analysis following rDNA damage. **A**. 100% stacked bar graph showing raw FBL abundance in MS analysis of nucleoplasm and nucleolar fractions from control and GRASP cells treated with 100 ng/ml doxycycline for 24h. R denotes “replicate”. The percentage of FBL levels in each compartment was calculated relative to the combined raw abundance of FBL in the nucleolus and nucleoplasm per condition **B**. Workflow of nucleoplasm and nucleolar proteome MS data analysis. Proteins with at least two valid values in at least one condition and coefficient of variation <1 between biological replicated were selected. The mean abundance of proteins for each nucleolar or nucleoplasm sample was normalized to the mean value of protein abundance across all 9 samples. The fold change in abundance of individual proteins between GRASP and control samples was calculated and significance in fold change in 3 biological replicates was measured using unpair t-test. Enriched or depleted proteins in nucleoli and nucleoplasm with p-value ≤ 0.05 were selected.

**Figure S3.**
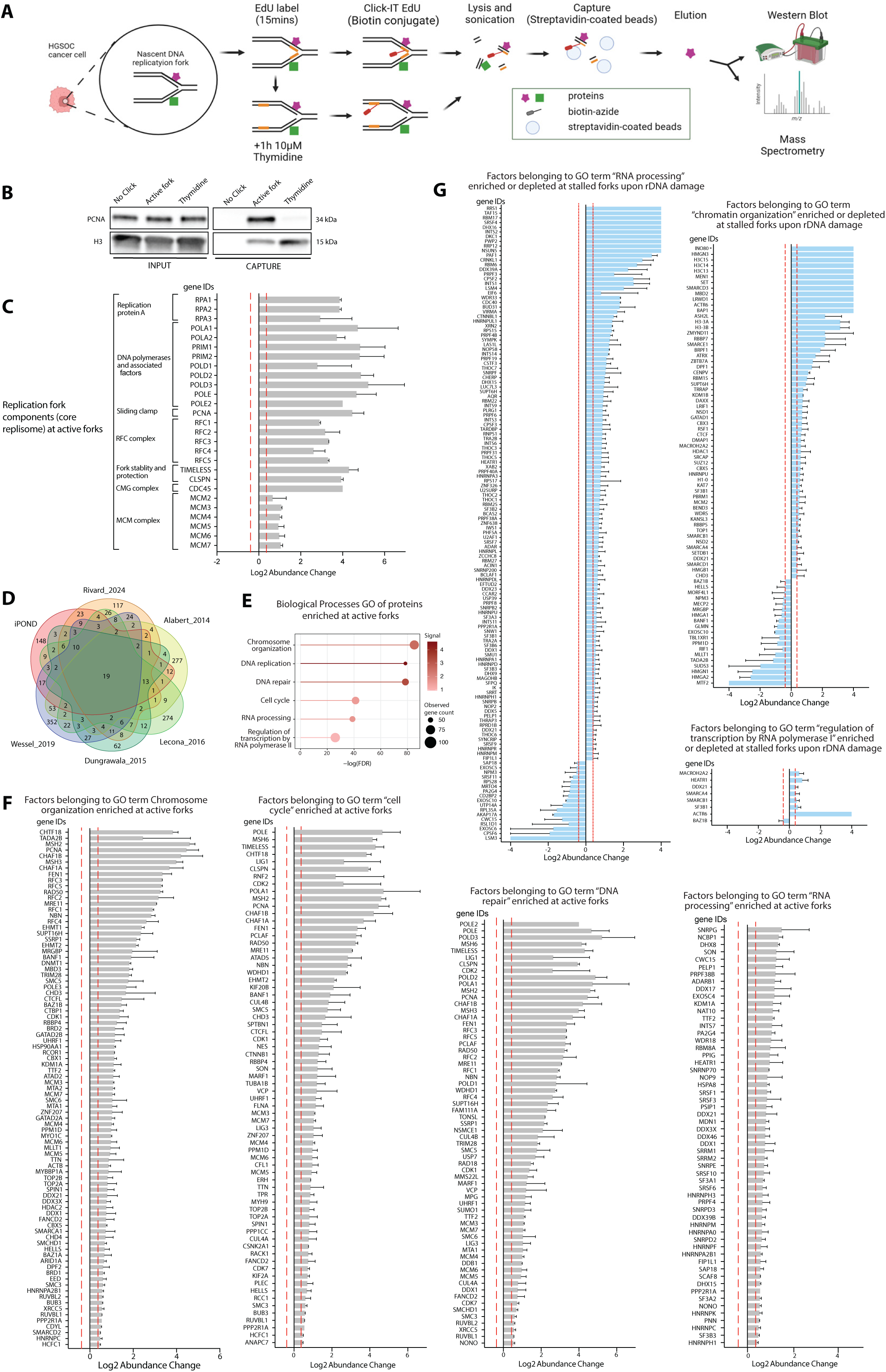
rDNA damage induces nucleolar replication stress. **A.** Schematic illustrating the aniPOND labelling and elution process. Active replication forks in control cells were collected 15 minutes after the EdU pulse. EdU-labeled cells treated with 10 mM thymidine for 1 hour as a control for chromatin-associated proteins. GRASP cells were treated with idoxycycline (100 ng/ml) for 24 hours followed by EdU labelling for 15 minutes for analysis of stalled replication sites. **B**. Western blot analysis of PCNA in aniPOND capture samples in OVCAR8 cells with Histone H3 serving as chromatin loading control. **C**. Log_2_ enrichment of core replisome proteins at active fork in control OVCAR8 cells treated with doxycycline (100 ng/ml) for 24 hours, compared with Thy chase chromatin controls. **D**. Comparison of enriched proteins at active forks in OVCAR8 cells across five replisome datasets ^17–21^. **E**. Lollipop plot of the significantly overrepresented biological processes of proteins enriched at active forks in control OVCAR8 cells. **F**. Log_2_ enrichment of factors involved in GO term chromosome organization, cell cycle, DNA repair and RNA processing at active fork in control OVCAR8 cells. **G**. Log_2_ change of factors involved in GO term RNA processing, chromatin organization and regulation of transcription by RNA Pol I at stalled replication forks in GRASP cells, compared with active replication forks in control OVCAR8 cells.

**Figure S4.**
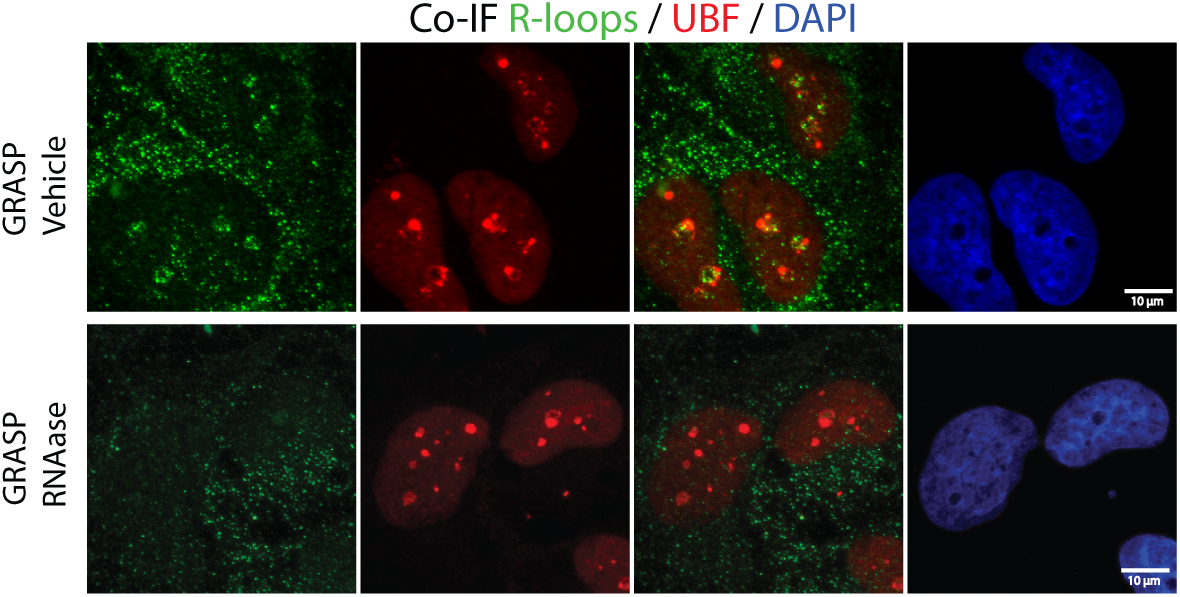
Pol II transcription-drives nucleolar R-loop formation Co-IF analysis of R-loops and UBF in GRASP 1 cells treated with doxycycline (100 ng/ml) for 24 hours, or GRASP 1 cells treated with doxycycline (100 ng/ml) for 24 hours and 100ul (50units/ml) RNase H for 3h leading to degradation of R-loops.

**Figure S5.**
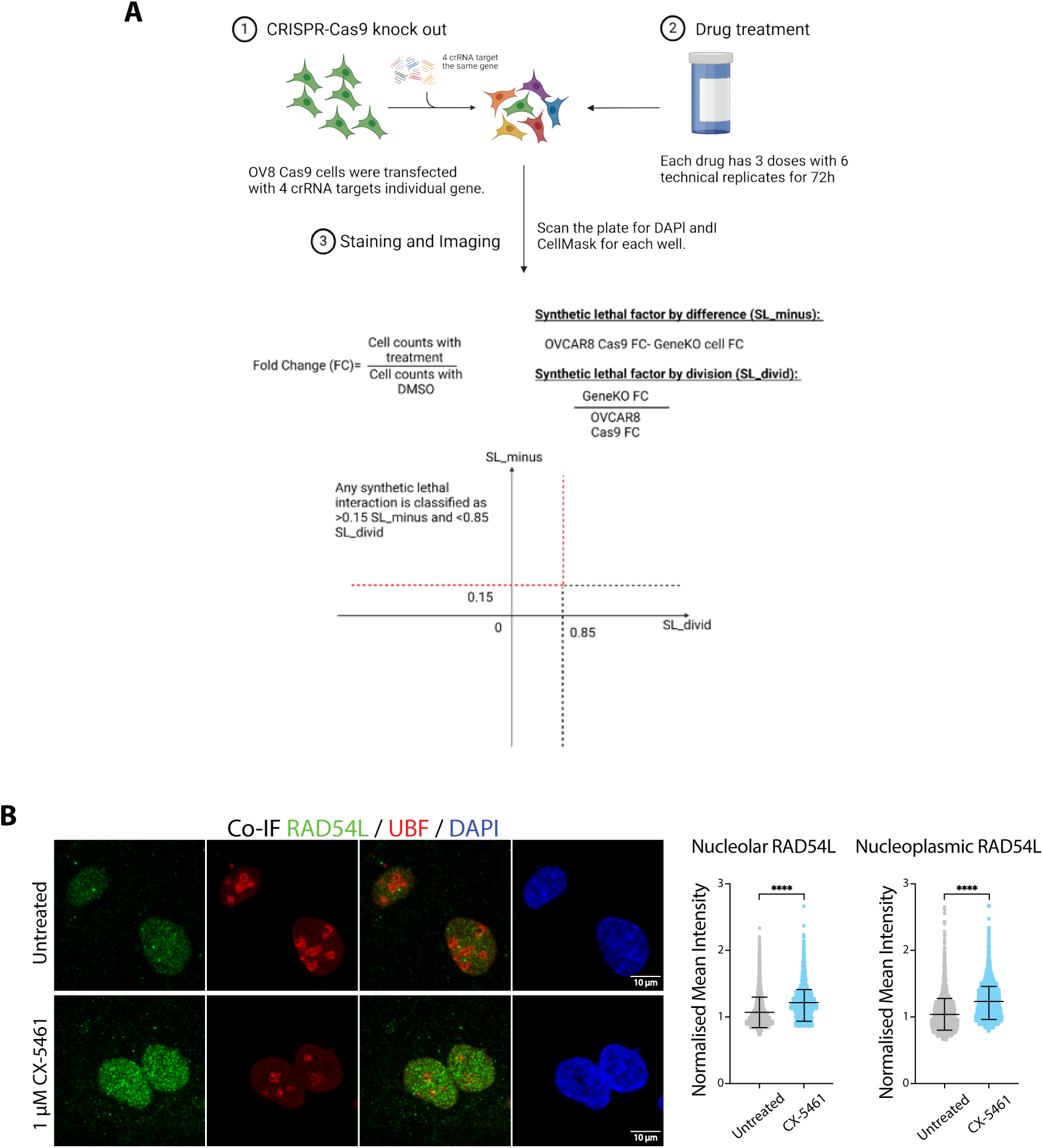
CRISPR-Cas9 screen of synthetic lethal interactions with DDR genes. **A.** Schematic representation of the workflow for the CRISPR-Cas9 Screen. OVCAR8 cells expressing Cas9 were reverse transfected with crRNAs per gene (Table S13). After 96 hours, cells were replated and treated with selected drugs for 72 hours, followed by staining with DAPI and imaging using Cellomics CX7 LZR (Thermo Fisher) at x10 magnification. Cell survival was assessed based on cell number determined from DAPI-stained cell counts. Significant synthetic lethal (SL) interactions were assessed by firstly calculating the fold change in cell proliferation in drug-treated KO cells relative to DMSO-treated gene KO cells. Two factors were then calculated: SL_minus and SL_divid, for each gene KO and drug treatment. SL_divid is the percentage change of gene KO after drug treatment compared to the change in parental OVCAR8 Cas9 with the same treatment. To avoid identifying gene KO with a strong effect on cell proliferation alone, the absolute difference was also calculated between gene KO and OVCAR9 Cas9 after drug treatment (SL_minus). Therefore, the synthetic lethal hits were selected as >0.15 of SL-minus and <0.85 of SL_divid. **B**. Co-IF analysis of RAD54L and NPM in OVCAR4 cells treated with or without 1 μM CX-5461 for 3h. Representative images from three independent experiments. Mean signal intensity for RAD54L was quantified using CellProfiler and normalized to the median value for control cells per experiment. Over 100 cells were analyzed per condition across three independent experiments. Error bars represent mean ± SD. Statistical analysis was conducted using an unpaired t-test; **** is p-value ≤ 0.0001.

## Notes

### Competing Interest Statement

The authors have declared no competing interest.

## References

1 Schneider, D. A. RNA polymerase I activity is regulated at multiple steps in the transcription cycle: recent insights into factors that influence transcription elongation. Gene 493, 176–184 (2012). 10.1016/j.gene.2011.08.006

2 Killen, M. W., Stults, D. M., Adachi, N., Hanakahi, L. & Pierce, A. J. Loss of Bloom syndrome protein destabilizes human gene cluster architecture. Hum Mol Genet 18, 3417–3428 (2009). 10.1093/hmg/ddp282

3 Xuan, J., Gitareja, K., Brajanovski, N. & Sanij, E. Harnessing the Nucleolar DNA Damage Response in Cancer Therapy. Genes (Basel) 12 (2021). 10.3390/genes12081156

4 Wang, M. & Lemos, B. Ribosomal DNA copy number amplification and loss in human cancers is linked to tumor genetic context, nucleolus activity, and proliferation. PLoS Genet 13, e1006994 (2017). 10.1371/journal.pgen.1006994

5 Warmerdam, D. O. & Wolthuis, R. M. F. Keeping ribosomal DNA intact: a repeating challenge. Chromosome Res 27, 57–72 (2019). 10.1007/s10577-018-9594-z

6 Lindstrom, M. S. et al. Nucleolus as an emerging hub in maintenance of genome stability and cancer pathogenesis. Oncogene 37, 2351–2366 (2018). 10.1038/s41388-017-0121-z

7 Goffova, I. & Fajkus, J. The rDNA Loci-Intersections of Replication, Transcription, and Repair Pathways. Int J Mol Sci 22 (2021). 10.3390/ijms22031302

8 van Sluis, M. & McStay, B. A localized nucleolar DNA damage response facilitates recruitment of the homology-directed repair machinery independent of cell cycle stage. Genes Dev 29, 1151–1163 (2015). 10.1101/gad.260703.115

9 Warmerdam, D. O., van den Berg, J. & Medema, R. H. Breaks in the 45S rDNA Lead to Recombination-Mediated Loss of Repeats. Cell Rep 14, 2519–2527 (2016). 10.1016/j.celrep.2016.02.048

10 Larsen, D. H. et al. The NBS1-Treacle complex controls ribosomal RNA transcription in response to DNA damage. Nat Cell Biol 16, 792–803 (2014). 10.1038/ncb3007

11 Mooser, C. et al. Treacle controls the nucleolar response to rDNA breaks via TOPBP1 recruitment and ATR activation. Nat Commun 11, 123 (2020). 10.1038/s41467-019-13981-x

12 Korsholm, L. M. et al. Double-strand breaks in ribosomal RNA genes activate a distinct signaling and chromatin response to facilitate nucleolar restructuring and repair. Nucleic Acids Res 47, 8019–8035 (2019). 10.1093/nar/gkz518

13 Kruhlak, M. et al. The ATM repair pathway inhibits RNA polymerase I transcription in response to chromosome breaks. Nature 447, 730–734 (2007). 10.1038/nature05842

14 Ciccia, A. et al. Treacher Collins syndrome TCOF1 protein cooperates with NBS1 in the DNA damage response. Proc Natl Acad Sci U S A 111, 18631–18636 (2014). 10.1073/pnas.1422488112

15 Boulon, S., Westman, B. J., Hutten, S., Boisvert, F. M. & Lamond, A. I. The nucleolus under stress. Mol Cell 40, 216–227 (2010). 10.1016/j.molcel.2010.09.024

16 Hamperl, S., Bocek, M. J., Saldivar, J. C., Swigut, T. & Cimprich, K. A. Transcription-Replication Conflict Orientation Modulates R-Loop Levels and Activates Distinct DNA Damage Responses. Cell 170, 774–786 e719 (2017). 10.1016/j.cell.2017.07.043

17 Rivard, R. S. et al. Improved detection of DNA replication fork-associated proteins. Cell Rep 43, 114178 (2024). 10.1016/j.celrep.2024.114178

18 Alabert, C. et al. Nascent chromatin capture proteomics determines chromatin dynamics during DNA replication and identifies unknown fork components. Nat Cell Biol 16, 281–293 (2014). 10.1038/ncb2918

19 Dungrawala, H. et al. The Replication Checkpoint Prevents Two Types of Fork Collapse without Regulating Replisome Stability. Mol Cell 59, 998–1010 (2015). 10.1016/j.molcel.2015.07.030

20 Lecona, E. et al. USP7 is a SUMO deubiquitinase essential for DNA replication. Nat Struct Mol Biol 23, 270–277 (2016). 10.1038/nsmb.3185

21 Wessel, S. R., Mohni, K. N., Luzwick, J. W., Dungrawala, H. & Cortez, D. Functional Analysis of the Replication Fork Proteome Identifies BET Proteins as PCNA Regulators. Cell Rep 28, 3497–3509 e3494 (2019). 10.1016/j.celrep.2019.08.051

22 Abraham, K. J. et al. Nucleolar RNA polymerase II drives ribosome biogenesis. Nature 585, 298–302 (2020). 10.1038/s41586-020-2497-0

23 Ketley, R. F. et al. DNA double-strand break-derived RNA drives TIRR/53BP1 complex dissociation. Cell Rep 41, 111526 (2022). 10.1016/j.celrep.2022.111526

24 Bywater, M. J. et al. Inhibition of RNA polymerase I as a therapeutic strategy to promote cancer-specific activation of p53. Cancer Cell 22, 51–65 (2012). 10.1016/j.ccr.2012.05.019

25 Peltonen, K. et al. A targeting modality for destruction of RNA polymerase I that possesses anticancer activity. Cancer Cell 25, 77–90 (2014). 10.1016/j.ccr.2013.12.009

26 Bossaert, M. et al. Transcription-associated topoisomerase 2alpha (TOP2A) activity is a major effector of cytotoxicity induced by G-quadruplex ligands. Elife 10 (2021). 10.7554/eLife.65184

27 Bruno, P. M. et al. The primary mechanism of cytotoxicity of the chemotherapeutic agent CX-5461 is topoisomerase II poisoning. Proc Natl Acad Sci U S A 117, 4053–4060 (2020). 10.1073/pnas.1921649117

28 Maclachlan, K. H. et al. Targeting the ribosome to treat multiple myeloma. Mol Ther Oncol 32, 200771 (2024). 10.1016/j.omton.2024.200771

29 Cameron, D. P. et al. CX-5461 Preferentially Induces Top2alpha-Dependent DNA Breaks at Ribosomal DNA Loci. Biomedicines 12 (2024). 10.3390/biomedicines12071514

30 Mason, J. M. et al. RAD54 family translocases counter genotoxic effects of RAD51 in human tumor cells. Nucleic Acids Res 43, 3180–3196 (2015). 10.1093/nar/gkv175

31 Uhrig, M. E. et al. Disparate requirements for RAD54L in replication fork reversal. Nucleic Acids Res 52, 12390–12404 (2024). 10.1093/nar/gkae828

32 Petropoulos, M. et al. Transcription-replication conflicts underlie sensitivity to PARP inhibitors. Nature 628, 433–441 (2024). 10.1038/s41586-024-07217-2

33 Kang, J. et al. Ribosomal proteins and human diseases: molecular mechanisms and targeted therapy. Signal Transduct Target Ther 6, 323 (2021). 10.1038/s41392-021-00728-8

34 Marszalek-Kruk, B. A. & Wojcicki, P. Identification of three novel TCOF1 mutations in patients with Treacher Collins Syndrome. Hum Genome Var 8, 36 (2021). 10.1038/s41439-021-00168-4

35 Sakai, D. & Trainor, P. A. Treacher Collins syndrome: unmasking the role of Tcof1/treacle. Int J Biochem Cell Biol 41, 1229–1232 (2009). 10.1016/j.biocel.2008.10.026

36 Hua, L., Yan, D., Wan, C. & Hu, B. Nucleolus and Nucleolar Stress: From Cell Fate Decision to Disease Development. Cells 11 (2022). 10.3390/cells11193017

37 Rawlinson, S. M. et al. Viral regulation of host cell biology by hijacking of the nucleolar DNA-damage response. Nat Commun 9, 3057 (2018). 10.1038/s41467-018-05354-7

38 Hiscox, J. A. RNA viruses: hijacking the dynamic nucleolus. Nat Rev Microbiol 5, 119–127 (2007). 10.1038/nrmicro1597

39 Burger, K. et al. Chemotherapeutic drugs inhibit ribosome biogenesis at various levels. J Biol Chem 285, 12416–12425 (2010). 10.1074/jbc.M109.074211

40 Sanij, E. et al. CX-5461 activates the DNA damage response and demonstrates therapeutic efficacy in high-grade serous ovarian cancer. Nat Commun 11, 2641 (2020). 10.1038/s41467-020-16393-4

41 Xu, H. et al. CX-5461 is a DNA G-quadruplex stabilizer with selective lethality in BRCA1/2 deficient tumours. Nat Commun 8, 14432 (2017). 10.1038/ncomms14432

42 Soberanis Pina PD, L. S., Han H, Shapiro G, Provencher DM, Rosen LS, Sardesai S, Taylor S, Cescon D, Alqaisi HA, Aparicio S, Sabantini P, Chao TI, Huang CE, Chen MC, Mahmud L, Ye XY, Bowering V, Oza AM. 631P Phase Ib expansion study of CX-5461 in patients with solid tumours and BRCA2 and/or PALB2 mutation. Developmental therapeutics 35 (2024).

43 Leung, K. H., Abou El Hassan, M. & Bremner, R. A rapid and efficient method to purify proteins at replication forks under native conditions. Biotechniques 55, 204–206 (2013). 10.2144/000114089

44 Wiest, N. E. & Tomkinson, A. E. Optimization of Native and Formaldehyde iPOND Techniques for Use in Suspension Cells. Methods Enzymol 591, 1–32 (2017). 10.1016/bs.mie.2017.03.001

45 Bensaddek, D., Nicolas, A. & Lamond, A. I. Quantitative Proteomic Analysis of the Human Nucleolus. Methods Mol Biol 1455, 249–262 (2016). 10.1007/978-1-4939-3792-9_20

46 Sanij, E. et al. UBF levels determine the number of active ribosomal RNA genes in mammals. J Cell Biol 183, 1259–1274 (2008). 10.1083/jcb.200805146

